# Redox regulated auto-processing controls delivery of an antibacterial cysteine peptidase toxin

**DOI:** 10.64898/2026.05.07.723447

**Authors:** Dinh Quan Nhan, Bonnie J. Cuthbert, Kaitlin A. Schroeder, Christine D. Hardy, Fernando Garza-Sánchez, Ian H. Matthews, Jack R. Kennaday, Michael S. Costello, Celia W. Goulding, Christopher S. Hayes

## Abstract

Contact-dependent growth inhibition (CDI) is a mechanism of inter-bacterial competition mediated by CdiA effectors, which deliver polymorphic C-terminal toxins (CT) into neighboring competitors. StbD from *Citrobacter rodentium* DBS100 is an unusual CdiA-like protein that carries a C-terminal cysteine peptidase toxin. Crystallography reveals that StbD-CT is composed of an N-terminal cytoplasm-entry domain connected to a C39 family peptidase by a flexible linker. The entry domain hijacks membrane-embedded YajC for translocation into the target-cell cytosol where the peptidase inactivates type II topoisomerases. Intoxication leads to a loss of DNA super-helicity, impaired chromosome segregation and cell filamentation. In addition to cleaving topoisomerases, StbD-CT exhibits auto-proteolytic processing under reducing conditions, and this activity is required for target cell intoxication. We propose that StbD-CT remains tethered to the cell periphery via interactions with YajC after delivery. Auto-processing releases the peptidase, enabling the domain to penetrate into the cell interior where it cleaves nucleoid-associated topoisomerases. Together, these findings identify a proteolytic effector that deactivates type II topoisomerases and reveal a redox regulatory strategy that coordinates toxin activation with intercellular delivery.

## Introduction

Gram-negative bacteria use a number of specialized secretion systems to translocate protein toxins directly into competing microbes (1). This phenomenon of ‘contact-dependent growth inhibition’ (CDI) was first described for *Escherichia coli* isolate EC93, which utilizes a type 5 secretion system (T5SS) to intoxicate neighboring bacteria (2). Subsequent studies have shown that other secretion systems, including type 6 (3, 4), type 4 (5) and type 1 (6), have been adapted for contact-dependent inter-bacterial competition, and it is now clear that cell-to-cell toxin delivery is a ubiquitous feature of prokaryotic ecology (1, 7). The originally discovered T5SS mode of CDI is mediated by a family of CdiB and CdiA two-partner secretion (TPS) proteins that are distributed widely across α-, β-, and γ-proteobacteria (8–11). CdiB is an outer-membrane localized transporter responsible for the export and assembly of CdiA effectors on the cell surface (12, 13). Upon recognition of specific receptors, CdiA transfers its C-terminal toxin region (CdiA-CT) into target bacteria to inhibit cell growth (14–18). Because toxins are also delivered into neighboring sibling cells, CDI^+^ strains produce cognate CdiI immunity proteins that neutralize CdiA-CT activity to prevent self-intoxication. CdiA-CT sequences are remarkably variable between bacteria, and different strains of the same species commonly deploy toxins with distinct activities (8, 19–21). In this manner, CDI systems provide a growth advantage against competitors in the environment, while neighboring isogenic sibling cells are protected from intoxication.

The mechanisms of CDI toxin delivery have been elucidated largely through studies in *E. coli* (22, 23), though the conserved domain architecture of CdiA across phyla suggests that these effectors use a common pathway (19, 24) (**Fig. 1A**). CdiA is first guided by the Sec machinery into the periplasm, where its N-terminal TPS transport domain is engaged by CdiB to initiate export across the outer membrane (13, 25). Based on studies with homologous FhaB-FhaC TPS proteins from *Bordetella* species, CdiA is thought to be transported through the lumen of CdiB as an unfolded polypeptide chain (13, 26, 27). As CdiA emerges from the cell, its filamentous hemagglutinin-1 (FHA-1) peptide repeat domain folds into a right-handed parallel β-helix (**Fig. 1B**) (28). FHA-1 domain assembly likely provides the driving force for export, and the resulting β-helical filament projects several hundred angstroms from the cell surface (**Fig. 1B**) (22). The distal end of the extracellular filament carries a receptor-binding domain (RBD), which recognizes specific ligands on the surface of target bacteria (15–18). Notably, CdiA export is halted by a small α-helical secretion-arrest (SA) domain shortly after the filament is assembled (**Figs. 1A** & **1B**). This export arrest imparts an unusual cell-surface topology, with the N-terminal half of CdiA forming an extracellular filament, while the C-terminal half is sequestered in the periplasm (**Fig. 1B**) (22). CdiA remains in this partially exported state until it engages its receptor, which triggers secretion to resume (**Fig. 1B**, steps 1 & 2). AlphaFold 3 modeling suggests that the newly secreted FHA-2 domain forms a coaxial β-helix extending from the tip of the FHA-1 filament (**Fig. 1B**, step 3) (29). The FHA-2 domain is also predicted to have long β-hairpins that form a large sheet emanating from the core β-helix. Because FHA-2 is required for toxin translocation into the target-cell periplasm (22), this domain presumably forms a β-barrel-like conduit across the outer membrane (**Fig. 1B**). Once transferred into the target-cell periplasm, the toxic CdiA-CT region is cleaved from the effector through pretoxin domain mediated autoproteolysis after a conserved VENN sequence motif (**Fig. 1B**, step 4) (22, 30). The released CdiA-CT fragment is composed of two domains: an N-terminal cytoplasm-entry domain that mediates cell import, and a C-terminal toxin with growth inhibition activity (**Fig. 1A**) (23). The entry domain hijacks inner membrane proteins to translocate into the target-cell cytosol through an ill-defined mechanism that requires the transmembrane proton gradient (23, 31–33). At least 29 types of entry domain have been identified in enterobacterial CdiA proteins (33), and effectors from *Burkholderia* and *Pseudomonas* species carry entry domains that exploit genus-specific membrane proteins (34–36). Thus, CDI toxin delivery is a complex multi-step process that requires protein translocation across four membranes.

**Figure 1.**
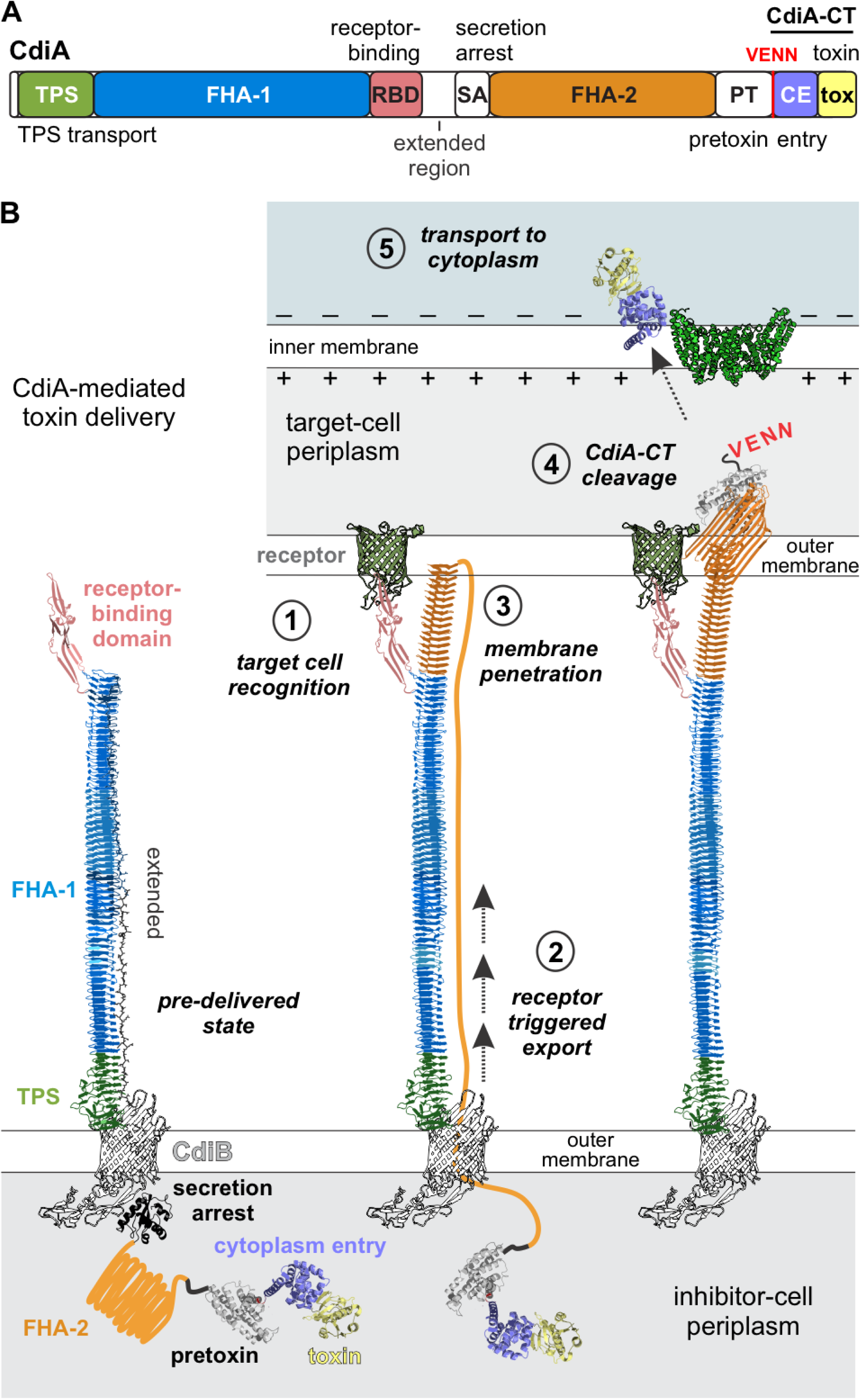
Model of CDI toxin delivery. **A**) Domain architecture of CdiA effector proteins. The CdiA-CT region is composed cytoplasm entry (CE) and C-terminal toxin domains. **B**) CdiA-mediated toxin delivery.

CDI toxins are extraordinarily polymorphic in sequence, yet every CdiA-CT characterized to date exhibits either nuclease or ionophore activity (8, 11, 19–21, 36–42). Informatic predictions suggest that a handful of CdiA effectors carry NAD glycohydrolase, deaminase and peptidase domains (43), but these biochemical activities have yet to be confirmed experimentally. Here, we characterize StbD from *Citrobacter rodentium* DBS100, which is an unusual CdiA-like protein with a C-terminal C39 cysteine peptidase domain. C39 peptidases were first identified as signal peptide processing domains within ATP-binding cassette (ABC) transporters that export microcins, lantibiotics and quorum-sensing peptides (44). Crystallography reveals that the C39 peptidase of StbD-CT most closely resembles quorum-sensing peptide processing domains from the ABC transporters of Gram-positive bacteria. StbD-CT intoxication results in target-cell filamentation and failure to segregate DNA nucleoids due to proteolytic inactivation of the GyrB and ParE subunits of DNA gyrase and topoisomerase IV, respectively. Although the C39 peptidase is specific for GyrB and ParE, it also catalyzes auto-cleavage of the peptide linker to the cytoplasm entry domain. Auto-cleavage is critical for toxin delivery during CDI, and we provide evidence that this redox-regulated processing enables the peptidase to penetrate into the target-cell cytosol.

## Results

### Citrobacter rodentium DBS100 encodes an unusual CdiA-like protein

*Citrobacter rodentium* DBS100 contains an unusual locus in which a type I fimbria gene cluster is fused to a *cdi* operon that lacks *cdiB* (**Fig. 2A**). As a result, the annotated *stbD* cistron encodes a CdiA-like effector with its N-terminal TPS transport domain replaced with a fimbria tip adhesin (NCBI: WP_012907078.1, **Figs. 2A** & **S1**). The operon retains chaperone (*stbB*) and usher (*stbC*) genes, but the major fimbria subunit *stbA* gene contains an in-frame amber stop codon at position 71 (**Fig. 2A**). We first tested whether this system mediates inter-cellular competition. We deleted the *stbD-CT* and *stbI* coding sequences to generate a DBS100 target strain that should be susceptible to intoxication by the C39 peptidase. However, the resulting Δ*stbD-CT/stbI* strain is not inhibited when co-cultured with wild-type DBS100 cells (**Fig. 2B**), suggesting that the *stbABCDI* operon is not functional or may not be expressed under laboratory conditions. To enforce expression, we introduced a rhamnose-inducible promoter upstream of *stbA* (**Fig. 2A**), but found that this strain also fails to inhibit immunity-less target bacteria (**Fig. 2B**). Given that CdiB and the TPS domain of CdiA are normally required for effector assembly (2, 13), we fused a rhamnose-inducible *cdiBÁ* construct from *E. coli* STEC_O31 in-frame with *stbD* to promote export (**Fig. 2A**). Immunoblot analysis showed DBS100 cells produce chimeric CdiA-StbD when induced with rhamnose (**Fig. 2C**, lane 6), yet this strain does not inhibit DBS100 Δ*stbD-CT/stbI* target bacteria in co-culture (**Fig. 2B**). Because CDI can be blocked by capsular polysaccharides found on wild bacteria (14, 17), we tested whether the inducible constructs inhibit domesticated *E. coli* strains that lack these carbohydrate layers. We first isolated the *stbABCDI* and *cdiBA-stbDI* gene clusters onto plasmid vectors for expression in *E. coli* MG1655. *E. coli* inhibitors that express *cdiBA-stbDI* outcompete target cells ∼100-fold during co-culture on agar, whereas induction of the wild-type *stbABCDI* operon confers no growth advantage (**Fig. 2D**). The CdiA-StbD chimera inhibits by virtue of the StbD-CT toxin because target bacteria are protected when they express the *stbI* immunity gene (**Fig. 2D**). Given that *E. coli* target cells are susceptible to StbD-CT intoxication, these data suggest that DBS100 may be intrinsically resistant to the chimeric CdiA-StbD effector.

**Figure 2.**
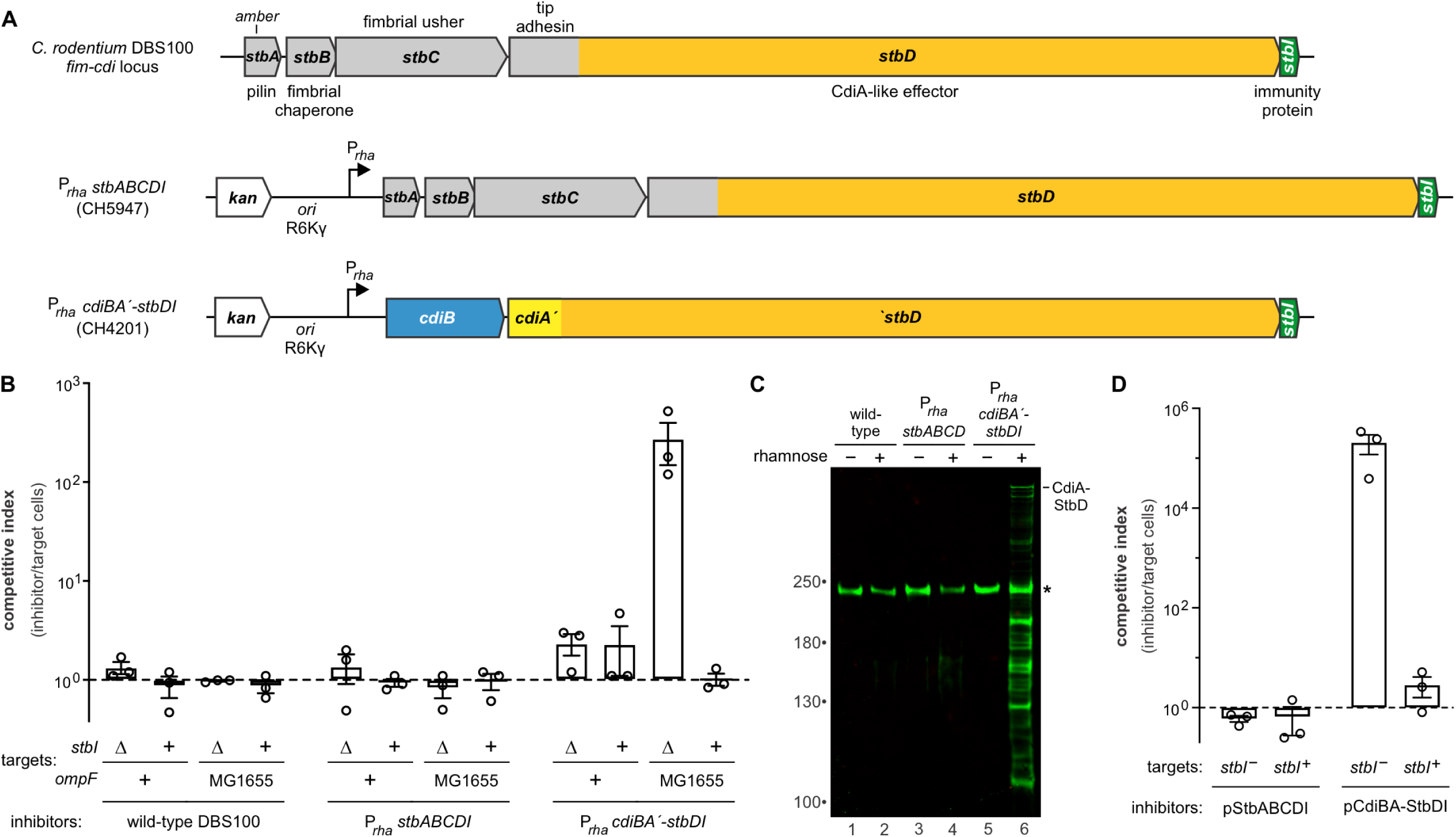
*Citrobacter rodentium* DBS100 encodes an unusual CdiA-like protein. **A**) *C. rodentium* DBS100 *stbABCDI* gene cluster. The *fim-cdi* operon is placed under the control of a rhamnose-inducible promoter (P*_rha_*) in strain CH5947. The *stbD* coding sequence is fused to *cdiBÁ* from *E. coli* STEC_O31 in strain CH4201 for rhamnose-inducible expression. **B**) *C. rodentium* growth competitions. *C. rodentium* inhibitor strains were mixed with DBS100 Δ*stbD-CT/stbI* target cells at a 1:1 ratio and cultured on solid media for 3 h at 37 °C. Viable inhibitor and target bacteria were enumerated and competitive indices calculated as the final ratio of inhibitor to targets divided by the initial ratio. Data are presented as the average ± standard error for three independent experiments. **C**) Immunoblot analysis of CdiA-StbD. Total urea-soluble protein was extracted from strains DBS100, CH5947 and CH4201 for immunoblotting with polyclonal antibodies raised against the N-terminal TPS domain of CdiA. The asterisk (*) indicates are cross-reacting protein produced in *C. rodentium*. **D**) *E. coli* growth competitions. *E. coli* inhibitor strains carrying the indicated plasmids were mixed with *E. coli* MG1655 target cells at a 1:1 ratio and cultured on solid media for 3 h at 37 °C. Viable inhibitor and target bacteria were enumerated and competitive indices calculated as the final ratio of inhibitor to targets divided by the initial ratio. Data are presented as the average ± standard error for three independent experiments.

### StbD exploits OmpF and YajC as receptors for toxin delivery

The receptor-binding domain (RBD) of StbD most closely resembles that of CdiA^EC536^ from *E. coli* 536 (**Fig. S1**), which recognizes OmpC and OmpF osmoporins on target bacteria (18, 45). Accordingly, we found that *E. coli* Δ*ompF* Δ*ompC* mutants are resistant to inhibition by CdiA-StbD (**Fig. 3A**). Complementation with plasmid-borne *ompF* and *ompC* revealed that OmpF is required for intoxication, whereas OmpC plays no role (**Fig. 3A**). AlphaFold 3 modeling predicts that the StbD RBD inserts a β-hairpin into the central lumen of OmpF (**Fig. 3B**) (46). The RBD is also predicted to interact with extracellular loops L1 and L3, and make additional contacts with loop L4 from an adjacent protomer in trimeric OmpF (**Figs. 3B**, **3C** & **3D**). OmpF^DBS100^ is ∼82% identical to OmpF^MG1655^ from *E. coli* MG1655, but there are significant differences in extracellular loops L4, L5 and L6 (**Fig. 3C**). AlphaFold 3 predicts van der Waals clashes between loop L4 of OmpF^DBS100^ and the StbD RBD, perhaps explaining why DBS100 is resistant to intoxication. Consistent with this hypothesis, complementation of the *E. coli ΔompF ΔompC* strain with *ompF*^DBS100^ leads to only a modest increase in toxin susceptibility (**Fig. 3A**), though OmpF^DBS100^ is produced at somewhat lower levels than OmpF^MG1655^ (**Fig. 3E**, lanes 2 & 4). We also found that expression of *ompF*^MG1655^ in DBS100 Δ*stbD-CT/stbI* target cells renders them sensitive to intoxication by the CdiA-StbD fusion (**Fig. 2B**). Thus, the 16-residue insertion within L4 of OmpF^DBS100^ appears to prevent RBD docking into the β-barrel.

**Figure 3.**
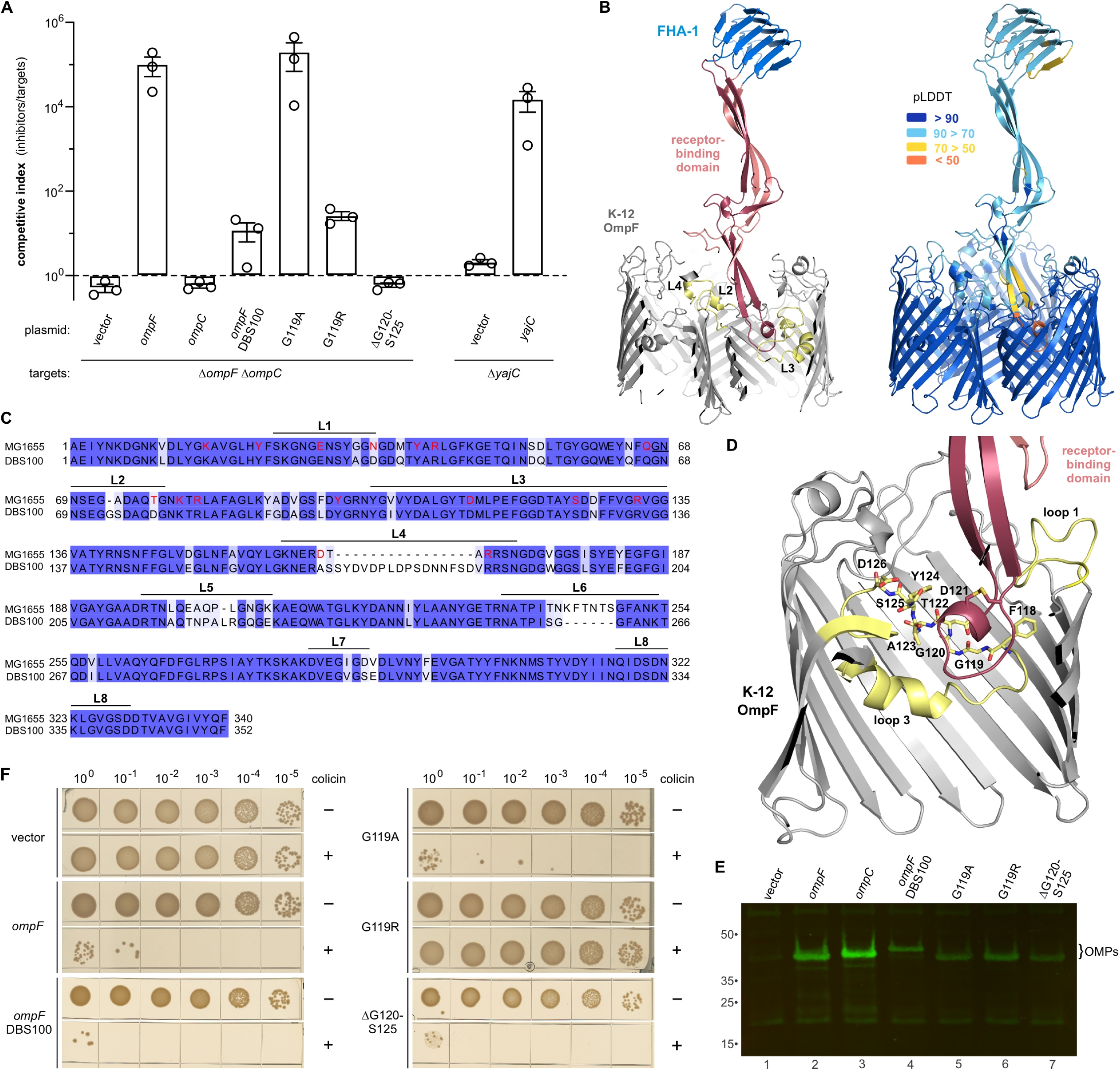
StbD exploits OmpF and YajC as receptors for toxin delivery. **A**) *E. coli* growth competitions. *E. coli* inhibitor cells that express *cdiA-stbD* were mixed with the indicated *E. coli* MG1655 target cells at a 1:1 ratio and cultured on solid media for 3 h at 37 °C. Viable inhibitor and target bacteria were enumerated and competitive indices calculated as the final ratio of inhibitor to targets divided by the initial ratio. Data are presented as the average ± standard error for three independent experiments. **B**) AlphaFold 3 model of the StbD receptor-binding domain (RBD) docked on trimeric OmpF. On the left, the RBD is shown in red, and interacting OmpF loops that are rendered in yellow. Predicted local distance difference test (pLDDT) values are indicated on the model to the right. **C**) Alignment of OmpF sequences from *E. coli* MG1655 and *C. rodentium* DBS100. Extracellular loops are indicated about the sequence, and residues predicted to make direct contact with the StbD receptor-binding domains are rendered in red font. **D**) AlphaFold 3 model of interactions with L3 of OmpF^MG1655^. **E**) Immunoblot analysis of OmpF variants. Total urea-soluble protein was extracted from *E. coli* Δ*ompF ΔompC* cells that express the indicated *omp* genes from a plasmid vector. Proteins were detected with polyclonal antibodies against *E. coli* OmpC. **F**) Colicin E5 killing of *E. coli* strains. *E. coli* Δ*ompF ΔompC* cells that express the indicated *omp* genes were treated with 100 nM purified colicin E5, then plated onto LB agar to enumerate surviving bacteria.

To interrogate the AlphaFold model further, we asked whether mutations that either constrict or enlarge the lumen of OmpF^MG1655^ affect receptor function. We first introduced Gly119Ala and Gly119Arg substitutions in L3, which are predicted to occlude the central pore progressively. The conservative Gly119Ala mutation has no discernable effect, but the Gly119Arg substitution confers significant resistance (**Fig. 3A**). Previous work has shown that the aperture of OmpF^MG1655^ can be enlarged by in-frame deletions in loop L3 (47, 48). We found that deletion of residues Gly120 through Ser125 from loop L3 (ΔGly120-Ser125) confers resistance comparable to that of the *ΔompF* null mutation (**Fig. 3A**). Given that OmpF^MG1655^ variants accumulate to somewhat lower levels than wild-type (**Fig. 3E**), we also tested whether the porins support colicin E5 toxin import. Colicin E5 uses OmpF as a portal to thread its unstructured N-terminus into the periplasm, where the toxin engages the Tol-Pal apparatus for internalization (49, 50). Thus, *E. coli* Δ*ompF ΔompC* cells are resistant to colicin E5, but become susceptible to intoxication when complemented with wild-type *ompF*^MG1655^ (**Fig. 3F**). Similarly, OmpF^DBS100^ supports colicin import (**Fig. 3F**), despite its poor recognition as a receptor by StbD (**Fig. 3A**). The OmpF^MG1655^ Gly119Ala variant also supports colicin import, but bacteria producing the Gly119Arg porin are resistant (**Fig. 3F**), consistent with occlusion of the central pore. Importantly, the ΔGly120-Ser125 variant supports colicin import to the same level as wild-type OmpF^MG1655^ (**Fig. 3F**). Given that OmpF ΔGly120-Ser125 provides a functional portal for colicin translocation, this porin must be properly assembled into the outer membrane. Together, these results suggest that the RBD makes direct contacts with OmpF loop L3 as predicted by AlphaFold modeling.

We next used a forward genetic approach to identify the inner-membrane receptor for StbD-CT, reasoning that mutants lacking this receptor should be resistant to intoxication. To avoid the isolation of resistant *ompF* mutants, we delivered StbD-CT into target bacteria using CdiA from *E. coli* EC93, which exploits the essential BamA protein as a cell-surface receptor (14, 15). After three cycles of selection, we obtained resistant clones from three of six *mariner* transposon mutant pools. Each isolate contained a unique insertion in the *yajC* gene, which encodes a small transmembrane protein associated with the general secretion machinery (51, 52). YajC is encoded in an operon with *secD* and *secF* genes, raising the possibility that resistance could be due to polar effects on *secD* or *secF* expression. However, a non-polar Δ*yajC* mutation confers resistance to StbD-CT intoxication, and target bacteria are resensitized when complemented with plasmid-borne *yajC* (**Fig. 3A**). Together, these data indicate that the entry domain of StbD-CT exploits YajC for translocation to the target-cell cytoplasm.

### The StbD-CT toxin undergoes redox regulated auto-processing

Having established that StbD-CT inhibits bacterial growth when delivered through the CDI pathway, we next examined the toxin’s biochemical activity. We cloned the *stbD-CT/stbI* coding sequences under the control of a T7 RNA polymerase driven promoter to over-produce the StbD-CT•StbI complex. A His_6_ tag was appended to the C-terminus of StbI to enable Ni^2+^-affinity purification of the toxin•immunity protein complex (**Fig. 4A**, lane 3). StbD-CT was isolated from the resin-bound complex by denaturation in 6 M guanidine-HCl (**Fig. 4A**, lane 1), followed by elution of StbI-His_6_ with imidazole (**Fig. 4A**, lane 2). Full-length StbD-CT is recovered after refolding by dialysis under oxidizing conditions (**Fig. 4A**, lane 1), but under reducing conditions the toxin is cleaved into fragments of ∼16 kDa (**Fig. 4A**, lane 4). Processing is not observed with the intact StbD-CT•StbI complex (**Fig. 4A**, lane 6), suggesting that the immunity protein blocks this activity. To test whether cysteine peptidase activity is required for cleavage, we substituted the predicted catalytic Cys183 residue (numbered from Val1 of the VENN motif) to alanine and appended an N-terminal His_6_ tag to facilitate purification. Wild-type StbD-CT carrying the N-terminal His_6_ tag is cleaved, but the Cys183Ala variant is not processed when refolded under reducing conditions (**Fig. 4B**, lanes 2 & 4). Mass spectrometry of the N-terminal fragment indicates that StbD-CT is cleaved between Gly158 and Ala159 in the predicted linker connecting the C39 peptidase to the N-terminal YajC-dependent entry domain (**Fig. 4C**). This cleavage occurs after a canonical diglycine (Gly157-Gly158) motif, and substitutions of these residues to serine block auto-processing (**Fig. 4D**, lanes 5 & 6). Given that reducing conditions promote cleavage, we hypothesized that Cys187 may control peptidase activity by forming a disulfide linkage with the catalytic Cys183 residue under oxidizing conditions. This model predicts that a Cys187Ser mutation should lead to constitutive auto-cleavage, but instead this substitution partially suppresses processing (**Fig. 4D**, lanes 7 & 8). Thus, the C39 peptidase domain is active and mediates auto-processing under reducing conditions.

**Figure 4.**
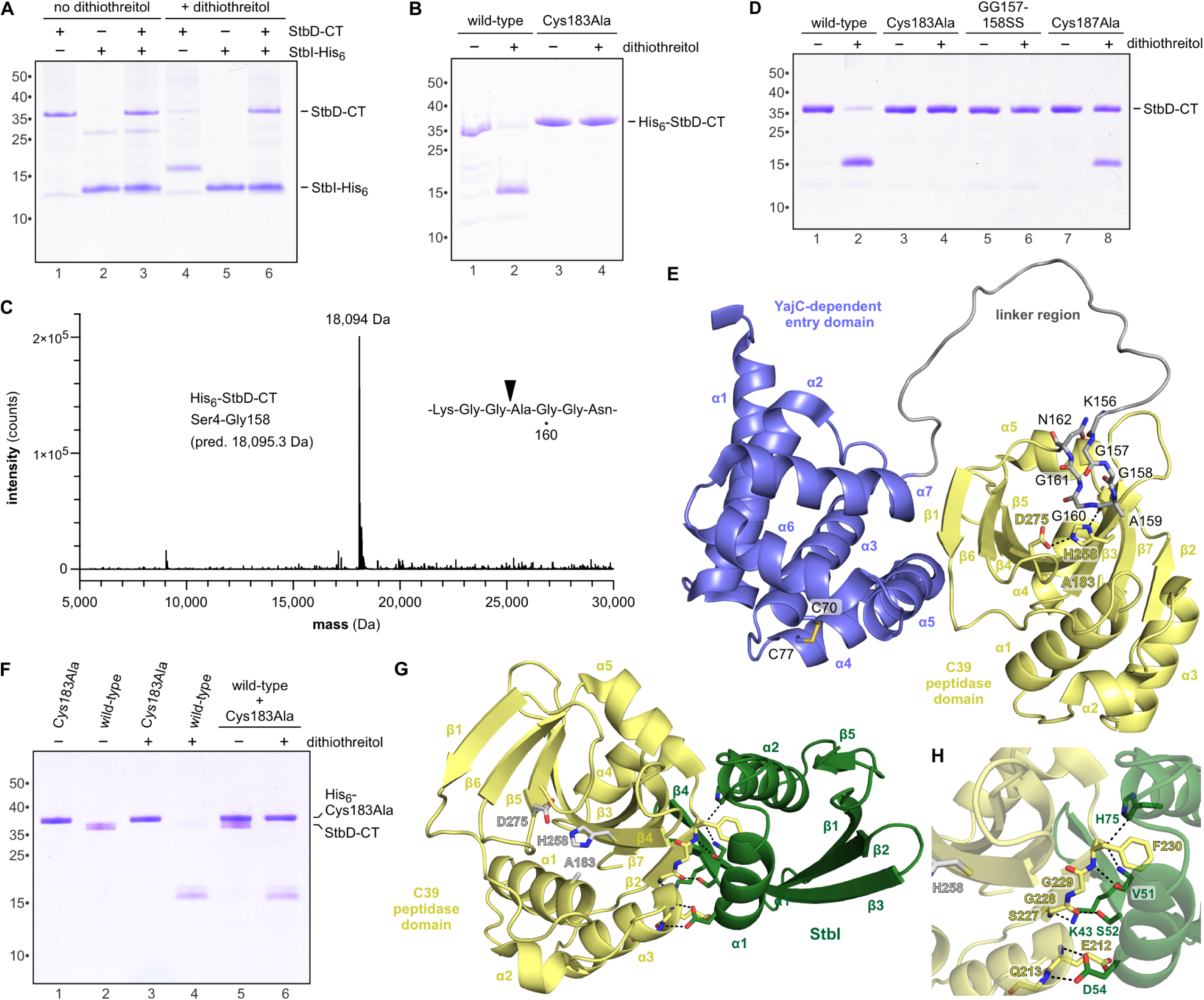
StbD-CT undergoes redox regulated auto-processing. **A**) StbD-CT auto-processing is induced with reducing agent. Purified StbD-CT and StbD-CT•StbI-His_6_ complex were dialyzed with and without dithiothreitol, then analyzed by SDS-PAGE. **B**) Auto-processing depends on C39 peptidase activity. **C**) Mass spectrometry reveals that StbD-CT auto-processing occurs after residue Gly158. **D**) Substitution of residues Gly157 and Gly158 abrogate C39 mediated auto-processing. **E**) Crystal structure of the inactive StbD-CT(C183A). StbD-CT is composed of an N-terminal YajC-dependent cytoplasmic entry domain and C-terminal C39 peptidase. **F**) Auto-processing occurs in *cis*. Active StbD-CT was mixed with His_6_-tagged StbD-CT(C183A) and incubated with and without dithiothreitol for SDS-PAGE analysis. **G**) Crystal structure of the C39•StbI complex. The C39 catalytic triad is rendered in grey, and polar interactions are indicated by dashed lines. **H**) C39 peptidase bonding network with StbI immunity protein.

### Crystal structures of StbD-CT and the C39 peptidase domain in complex StbI

To gain insight into auto-processing, we determined the crystal structure of inactive StbD-CT(C183A) to 1.9 Å resolution (**Fig. 4E** & **Table 1**). StbD-CT is composed of N-terminal cytoplasm entry and C-terminal C39 peptidase domains, which are connected by a flexible linker extending from Pro135 to Val165 (**Fig. 4E**). The asymmetric unit contains four StbD-CT molecules packed as a dimer of A/B and C/D dimers. The individual protomers align well with ∼0.7 Å root mean square deviation (rmsd) over ∼300 residues. The YajC-dependent entry domain is composed of seven α-helices with a partially formed disulfide [78% chain A, 78% chain B, 100% chain C, 80% chain D] linking Cys70 and Cys77 (**Fig. 4E**). The C39 peptidase domain is composed of a central seven-stranded antiparallel β-sheet (β1-β6-β5-β4-β3-β7-β2) decorated by helices α1, α2 and α3 on one side and helices α4 and α5 on the other (**Fig. 4E**). The C39 domain is similar in structure to the N-terminal domains of ComA from *Streptococcus mutans* (PDB:3K8U, 2.6 Å rmsd) (53) and LahT from *Lachnospiraceae* (PDB:6MPZ, 3.3 Å rmsd) (54). Superposition of StbD-CT onto ComA suggests that toxin residues Cys183 (mutated to Ala183 in the structure), His258 and Asp275 comprise the peptidase active site (53). The side-chain of His258 forms a hydrogen bond with the carbonyl of Ala159 in the cleavage site (**Fig. 4E**), suggesting that the inactive toxin is poised for processing in its crystal form. The structure also suggests that auto-processing is an intra-molecular reaction. To test this hypothesis, we asked whether the linker region of inactive StbD-CT(C183A) can be cleaved by the wild-type peptidase toxin *in vitro*. Although wild-type StbD-CT is processed in these reactions, the inactive StbD-CT(C183A) construct remains intact (**Fig. 4F**, lane 6), consistent with an obligate intramolecular cleavage reaction.

**Table 1.**
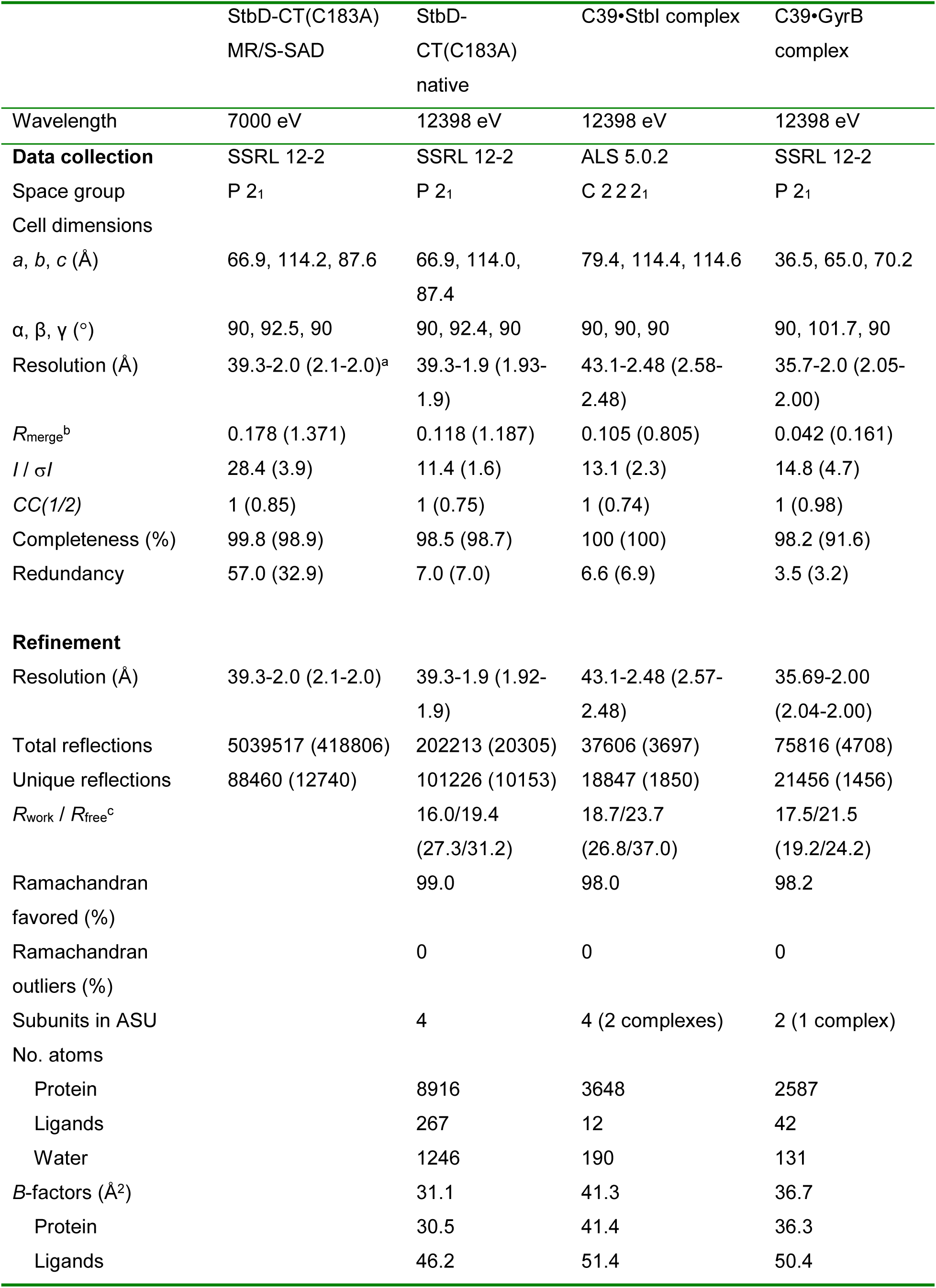
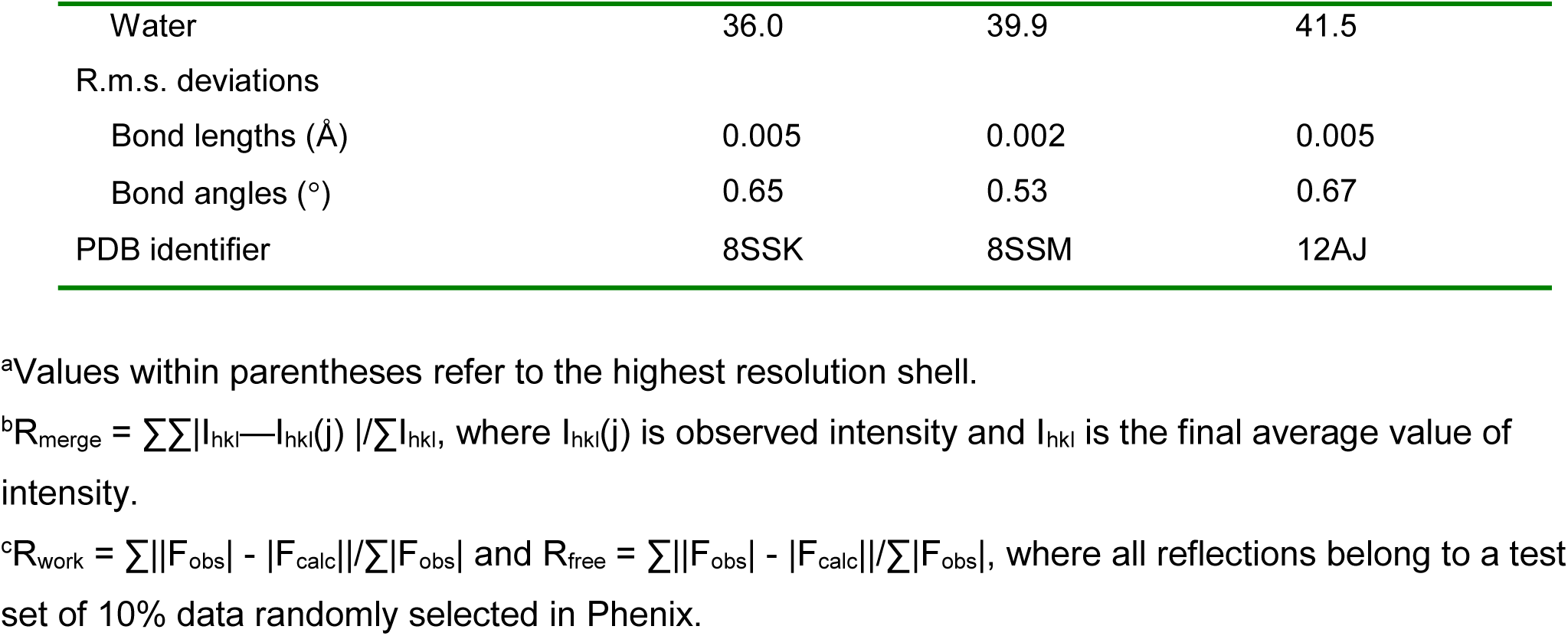
Crystallography data collection and refinement statistics.

We also solved the crystal structure of the inactive C39 peptidase domain (residues Ala159 – Gln301) in complex with the immunity protein to 2.5 Å resolution (**Fig. 4G** & **Table 1**). The C39•StbI complex crystallized with two heterodimers (chains A/D and B/C) in the asymmetric unit. The complexes are very similar with 0.71 Å rmsd over 232 total residues. StbI forms a curved five-stranded antiparallel β-sheet (β4-β3-β2-β1-β5) with two helices along the toxin•immunity interface (**Fig. 4G**). StbI cradles the peptidase along the opposite side of the active site through a network of 8-9 hydrogen bonds (**Figs. 4G** & **4H**, **Table S1**). StbI helix α1 interacts with peptidase helix α3, with the sidechain of StbI residue Asp54 forming hydrogen bonds to the backbone amides of Glu212 and Gln213 in the peptidase (**Figs. 4G** & **4H**). Immunity protein residues Lys43 and Ser52 make contact with peptidase residues Ser227 and Gly228, respectively (**Fig. 4H**). There are also backbone interactions between peptidase Phe230 and StbI residues Val51 and His75 (**Fig. 4H**). One complex has an additional hydrogen bond between Arg216 of the peptidase and StbI residue Asp54. This hydrogen-bonding network links peptidase strand β2 with β4 of StbI to create a 12-stranded mixed β-sheet across the complex.

### StbD-CT inactivates the ATPase subunits of DNA gyrase and topoisomerase IV

To gain insight into peptidase substrates, we examined target bacteria intoxicated by inhibitor cells that deploy StbD-CT. Fluorescence microscopy revealed elongated, filamentous target cells, some of which contained multiple unsegregated nucleoids (**Fig. 5A**). Filamentation was not observed when target bacteria express the *stbI* immunity gene (**Fig. 5A**), indicating that the morphological changes are a consequence of StbD-CT activity. The StbD-CT intoxication phenotype is reminiscent of the effects observed with quinolone and aminocoumarin antibiotics, which inhibit the activity of type II topoisomerases (55, 56). Indeed, similar morphological changes occur when target bacteria are treated with either nalidixic acid (quinolone) or novobiocin (aminocoumarin) (**Fig. 5A**). Further, intoxication leads to a loss of plasmid supercoiling in target bacteria (**Fig. 5B**), consistent with the inhibition of DNA gyrase activity. These observations suggest that topoisomerase IV and DNA gyrase are inhibited during intoxication.

**Figure 5.**
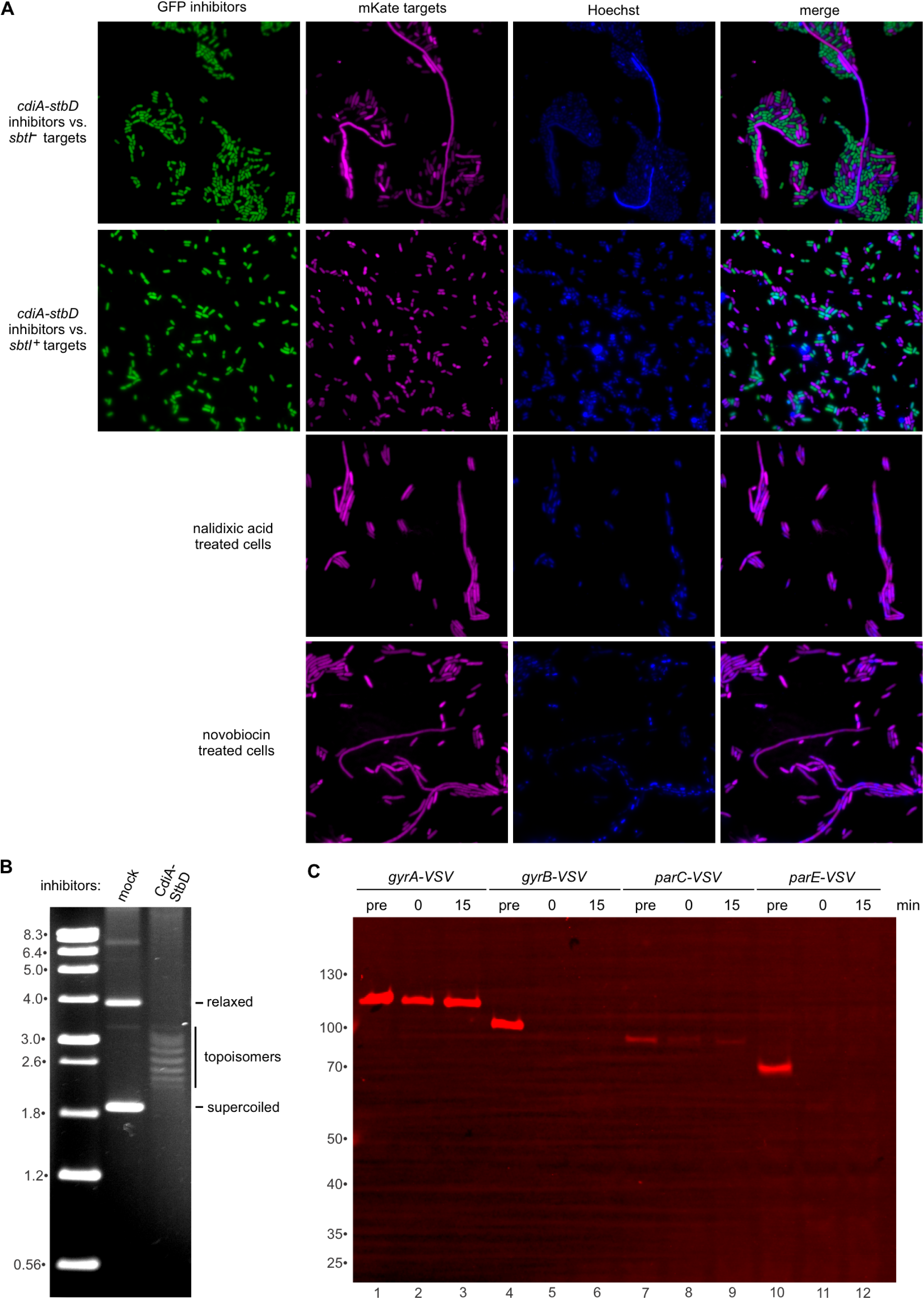
StbD-CT inactivates DNA gyrase and topoisomerase IV. **A**) Fluorescence microscopy of competition co-cultures. GFP labeled inhibitors were co-cultured with mKate labeled target bacteria in LB media, then spotted onto agarose pads for microscopic visualization. **B**) Plasmid supercoiling is perturbed in StbD-CT intoxicated bacteria. Inhibitors were mixed with targets that harbor pBluescript in LB medium for 30 min, then plasmid DNA was isolated for analysis on a chloroquine agarose gel. **C**) StbD-CT intoxication leads to degradation of GyrB and ParE. The VSV-G peptide epitope was appended to subunits of DNA gyrase and topoisomerase IV to enable immunoblot tracking in target bacteria.

DNA gyrase and topoisomerase IV are homologous hetero-tetrameric A_2_B_2_ enzymes that modulate chromosomal topology and DNA super-helicity (57). The A subunits (GyrA and ParC) use active-site Tyr residues to generate reversible double-strand breaks in gate segment (G-segment) DNA. The B subunits (GyrB and ParE) contain N-terminal ATPase domains that function as dynamic gates (N-gates) to capture duplex transport segment (T-segment) DNA for passage across the cleaved G-segment (58). To test whether type II topoisomerases are peptidase substrates, we appended C-terminal VSV-G epitope tags to each subunit at their native chromosomal loci, enabling the endogenous proteins to be monitored by immunoblot analysis. This approach revealed that GyrB-VSV and ParE-VSV are degraded when StbD-CT is delivered into target bacteria, while GyrA-VSV and ParC-VSV remain intact (**Fig. 5C**). Together, these data suggest that the C39 peptidase toxin inhibits target-cell growth by inactivating GyrB and ParE.

### StbD-CT cleaves a conserved Gly-Gly motif in GyrB and ParE

To ascertain whether GyrB and ParE are indeed substrates for StbD-CT, we purified His_6_-tagged versions of each topoisomerase subunit for *in vitro* peptidase assays. StbD-CT cleaves GyrB readily, whereas ParE is processed less efficiently (**Fig. 6A**, lanes 1 & 3). Importantly, cleavage is not observed when the peptidase reactions are supplemented with purified StbI immunity protein (**Fig. 6A**, lanes 2 & 4). GyrB is also cleaved efficiently in the context of the intact DNA gyrase complex containing GyrA (**Fig. 6B**, lane 2). Inclusion of plasmid DNA in the latter reaction has little effect on GyrB cleavage (**Fig. 6B**, lane 3). We found that substitutions of StbD-CT catalytic residues Cys183, His258 and Asp275 abrogate *in vitro* peptidase activity (**Fig. 6C**, lanes 6, 8 & 9), whereas mutations that interfere with StbD-CT auto-processing have no effect (**Fig. 6C**, lanes 4, 5 & 7). The large digestion fragments of GyrB and ParE retain C-terminal His_6_ tags, indicating that the peptidase likely cleaves within the N-terminal ATPase domain of each subunit. We purified ATPase domains from GyrB and ParE for *in vitro* cleavage assays and again found that GyrB is cleaved more efficiently than ParE (**Fig. 6D**, lanes 4 & 9). Mass spectrometry of the C-terminal GyrB-His_6_ fragment revealed cleavage between residues Gly101 and Gly102 (**Fig. 6E**). We note that ParE contains the same diglycine motif corresponding to residues Gly97 and Gly98 within a conserved His-Ala-Gly-Gly-Lys motif (**Fig. S2**). Although individual Gly101Ser and Gly102Ser substitutions do not block cleavage, a combination of both mutations prevents digestion of the ATPase domain (**Fig. 6D**, lanes 5-7). We also examined serine substitutions at another diglycine sequence (Gly113-Gly114) in GyrB, but these mutations have no discernable effect on toxin-mediated cleavage (**Fig. 6D**, lane 8).

**Figure 6.**
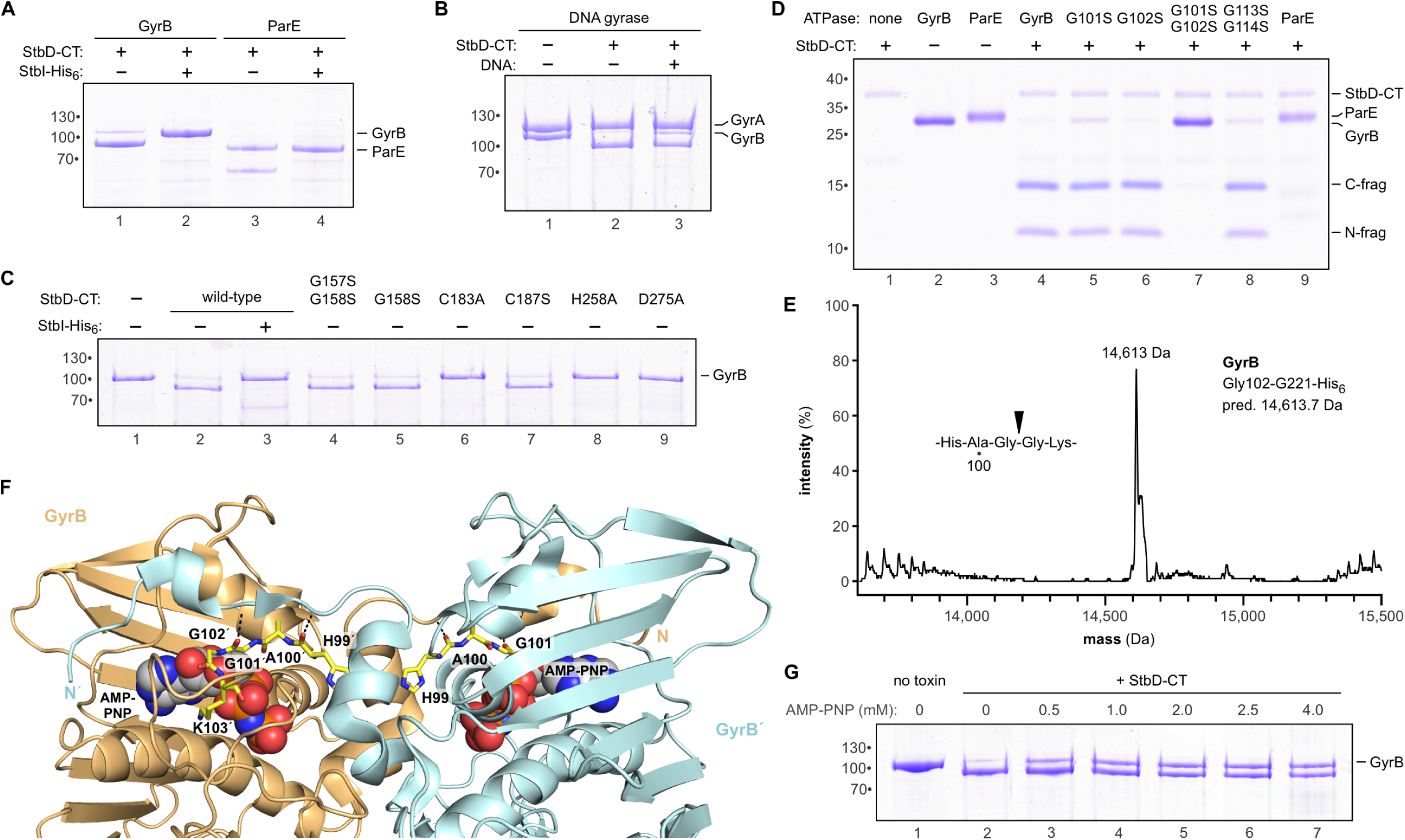
StbD-CT cleaves a conserved Gly-Gly motif in GyrB and ParE. **A**) Purified GyrB and ParE are cleaved by StbD-CT *in vitro*. **B**) GyrB is cleaved by StbD-CT in the context of assembled DNA gyrase. **C**) Peptidase activities of StbD-CT variants. The indicated StbD-CT variants were incubated with purified GyrB for 30 min at 37 °C, then analyzed by SDS-PAGE. **D**) StbD-CT cleaves the N-terminal ATPase domain of GyrB. The indicated variants of the GyrB ATPase domain were incubated with StbD-CT for 30 min at 37 °C, then analyzed by SDS-PAGE. **E**) Mass spectrometry of treated GyrB ATPase domain reveals cleavage after residue Gly101. **F**) Crystal structure of *E. coli* GyrB ATPase domain. GyrB is dimeric with flexible N-terminal arms extending to secure the ATP-binding sites of the opposite protomers. Model rendered from PDB:1EI1 (61). **G**) Purified GyrB was incubated with increasing concentrations of AMP-PNP, then treated with StbD-CT for 30 min at 37 °C before analysis by SDS-PAGE.

Residues Gly101 and Gly102 are presumably critical for GyrB ATPase activity, because they make direct contacts with the nucleotide (**Fig. 6F**) (59, 60). However, in the ATP-bound state, the scissile peptide bond is inaccessible because it is covered by the extended N-terminal arm of the other GyrB protomer (**Fig. 6F**) (59, 61). These interactions secure the ATP-binding pocket and are important for dimerization dependent closure of the N-gate (61). These observations suggest that GyrB should be resistant to cleavage in the ATP-bound state. Consistent with this model, we found that non-hydrolyzable AMP-PNP partially suppresses StbD-CT mediated cleavage of GyrB (**Fig. 6G**). These findings suggest that the C39 peptidase cleaves GyrB only in the apo state.

### Interaction between StbD-CT and GyrB

To explore how the toxin recognizes its substrate, we determined the crystal structure of the inactive Cys183Ala peptidase domain in complex with the GyrB ATPase domain to 2.0 Å resolution (**Table 1**). Although GyrB residues Asp106 - Gly114 are not resolved in the final model, there are clear conformation changes in the His-Ala-Gly-Gly-Lys motif compared to the structure of nucleotide-bound dimeric GyrB (**Fig. 7A**) (61). Lys103 of GyrB is displaced by ∼11 Å to form a salt-bridge with peptidase active-site residue Asp275, and the scissile Gly101-Gly102 peptide is positioned for nucleophilic attack by the catalytic Ala(Cys)183 residue (**Fig. 7A**). A series of main-chain hydrogen bonds stabilize the displaced GyrB loop (**Table S2**), with toxin residue Gly257 interacting with His99, Gly101 and Gly102 of the GyrB ATPase domain (**Fig. 7B**, top). Other contacts are made by peptidase residues Ala(Cys)183, Lys209 and the amide side-chain of Gln177 (**Fig. 7B**, top). In addition, the N-terminal arm of GyrB wraps around the peptidase on the side opposite from the catalytic center (**Fig. 7B**, bottom). These latter interactions are again dominated by main-chain hydrogen bonds (**Table S2**), though the side-chains of GyrB Arg22 and Tyr26 make prominent contacts (**Fig. 7B**, bottom). Notably, the structure of the C39•StbI complex indicates that the immunity protein should block these interactions with GyrB (**Figs. 7B**, bottom & **7C**), explaining how the peptidase is neutralized though its active site is exposed. All of the interacting GyrB residues are conserved in ParE (**Fig. S2**), and AlphaFold 3 modeling suggests that the peptidase binds ParE in a similar manner (**Figs. S2** & **S3**). Notably, the latter computational model is obtained only in the absence of nucleotide. When ATP is included, the N-terminal arms of ParE remain bound to the neighboring protomers, and AlphaFold 3 docks the C39 domain at various positions on the C-terminal TOPRIM domain of the topoisomerase.

**Figure 7.**
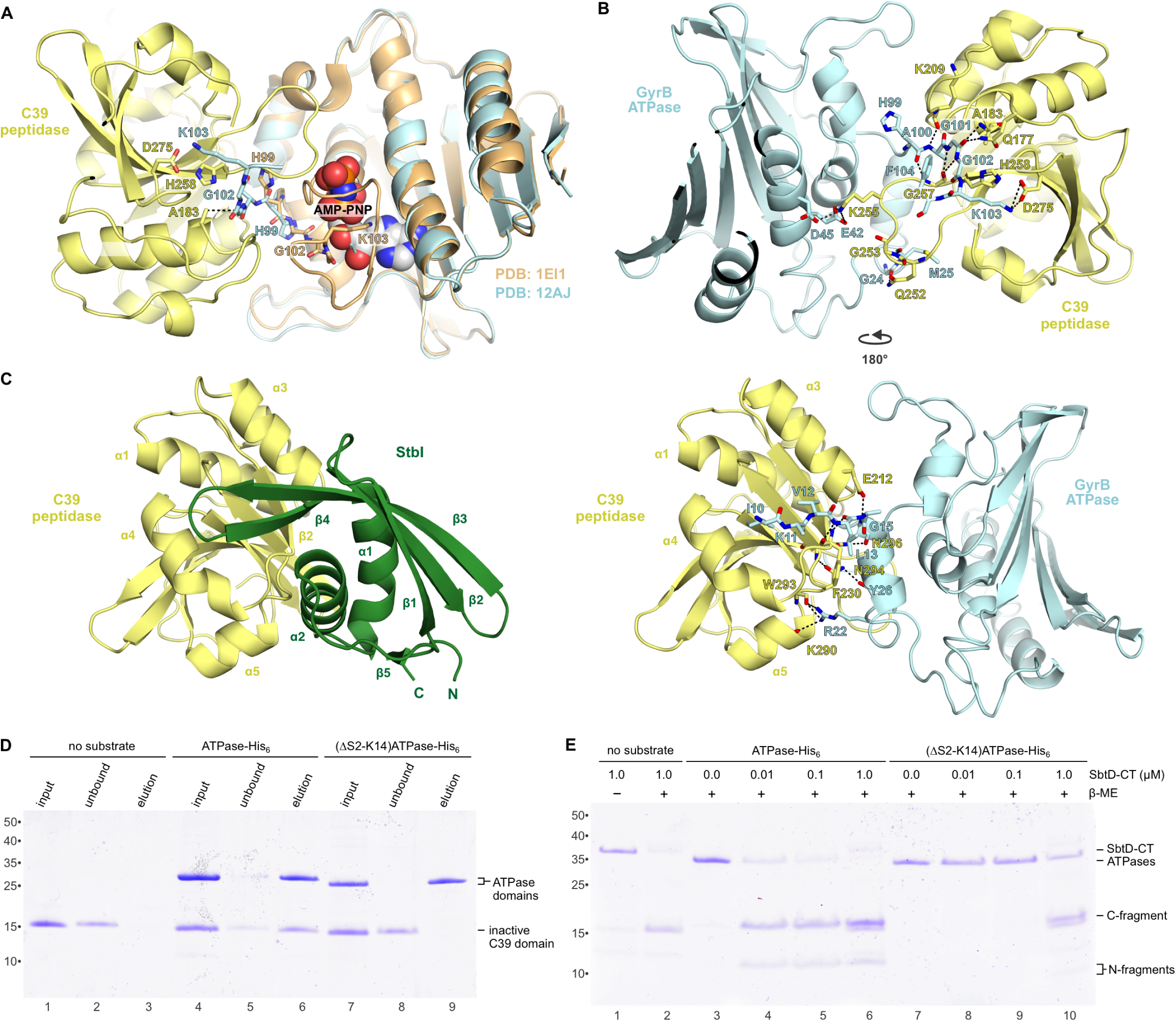
Interaction between the C39 peptidase and GyrB ATPase domain. **A**) Superposition of the C39•GyrB ATPase structure (PDB:12AJ) onto the GyrB ATPase domain from PDB:1EI1. Residues Gly101, Gly102 and Lys103 are displaced ∼11 Å from positions in the nucleotide-bound domain to interact with the peptidase active site. **B**) Structure of the C39•GyrB complex. The hydrogen-bonding network linking the C39 peptidase and GyrB ATPase domain is indicated by dashed lines. **C**) C39•StbI complex shown from the same perspective as the bottom view of panel B to illustrate how the immunity protein prevents interaction with GyrB. **D**) The N-terminal arm of GyrB contributes to high-affinity binding to C39. **E**) GyrB ATPase domains were incubated with increasing concentrations of StbD-CT.

The crystal structure of the C39•GyrB complex suggests that the N-terminal arm of the ATPase domain may be required for stable interaction with the toxin. To test this hypothesis, we used affinity co-purification to monitor the binding of inactive C39(C183A) peptidase to the GyrB ATPase domain. Untagged peptidase alone does not bind Ni^2+^-agarose resin, but it co-elutes with the His_6_-tagged ATPase domain during Ni^2+^-affinity chromatography (**Fig. 7D**, lanes 3 & 6). The N-terminal arm of GyrB contributes to the binding interaction because inactive peptidase does not co-purify with an ATPase domain construct lacking residues Ser2 - Lys14 (**Fig. 7D**, lane 9). *In vitro* peptidase assays revealed that the (ΔS2-K14)ATPase domain is also cleaved less efficiently by active StbD-CT toxin. The wild-type ATPase domain is digested in reactions containing 10 nM StbD-CT (**Fig. 7E**, lane 4), whereas cleavage of the truncated (ΔS2-K14)ATPase requires 100-fold more toxin (**Fig. 7E**, lane 10). Collectively, these results indicate that the toxin only acts on apo-GyrB and that the flexible N-terminal arms must be displaced to reveal the scissile peptide.

### StbD-CT auto-processing promotes target cell intoxication

Previous work has shown that CdiA-CT nucleases are released from full-length CdiA after transfer into the target-cell periplasm (22, 33). This auto-processing reaction is catalyzed by the CdiA pretoxin domain, which likely utilizes an asparagine lyase mechanism to cleave after the VENN motif (**Fig. 1B**, step 4) (22, 30). Given that CDI toxin delivery is already known to require auto-proteolytic release, the significance of C39 peptidase mediated auto-processing is not clear. We introduced Gly157Ser/Gly158Ser substitutions into full-length CdiA-StbD to block C39 auto-processing, then examined the growth inhibition activity in competition co-cultures. These mutations nearly abrogate target-cell killing, reducing the competitive fitness to that of mock inhibitors that carry empty vector plasmids (**Fig. 8A**). Mutation of the peptidase’s non-catalytic Cys187 residue, which is required for efficient auto-processing *in vitro* (see **Fig. 4D**, lane 8) has a more modest effect on growth inhibition activity (**Fig. 8A**). Notably, these mutations have no effect on peptidase activity with purified GyrB substrate *in vitro* (see **Fig. 6C**, lanes 4 & 7). These data indicate that auto-cleavage between the entry and peptidase domains is critical for target cell intoxication.

**Figure 8.**
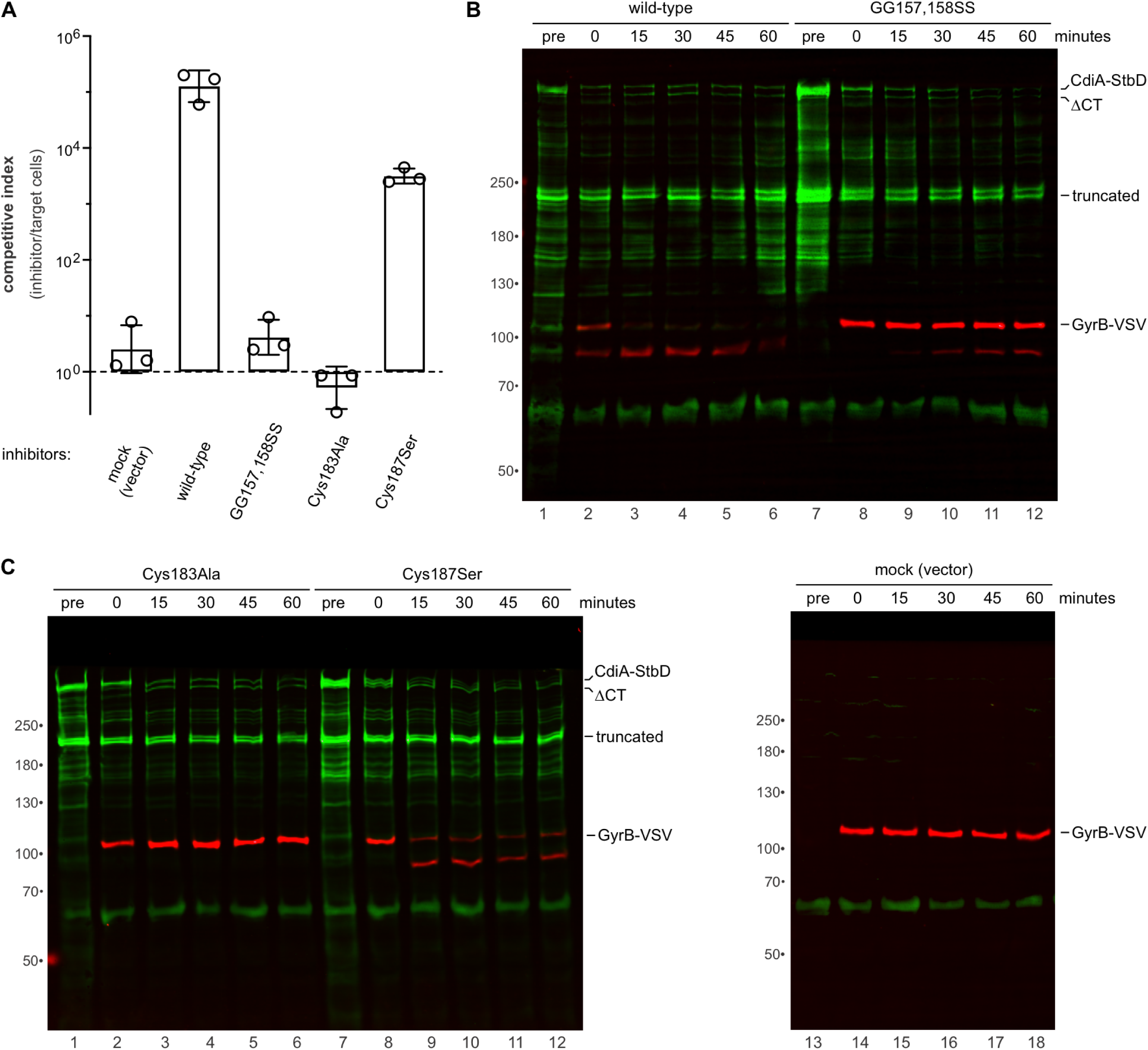
StbD-CT auto-processing promotes target cell intoxication. **A**) *E. coli* growth competitions. *E. coli* inhibitor cells that deliver the indicated StbD-CT variants were mixed with *E. coli* MG1655 target cells at a 1:1 ratio and cultured on solid media for 3 h at 37 °C. Viable inhibitor and target bacteria were enumerated and competitive indices calculated as the final ratio of inhibitor to targets divided by the initial ratio. Data are presented as the average ± standard error for three independent experiments. **B** & **C**) Immunoblot analysis of StbD-CT delivery. Inhibitor cells that deliver the indicated StbD-CT variants were treated with chloramphenicol for 30 min, then mixed at 1:2 ratio with chloramphenicol-resistant *E. coli gyrB-VSV* target bacteria. Samples were taken at each time point for immunoblot analysis with antibodies to CdiA (green) and the VSV epitope (red).

Finally, we used immunoblotting to examine StbD-CT processing during delivery. To maintain CdiA-StbD in the pre-delivered state, we expressed the effector protein in *E. coli* Δ*ompF* cells, which lack the receptor for StbD. Under these conditions, CdiA-StbD is produced as the full-length effector and a major truncated form (**Fig. 8B**, lane 1). Previous studies have shown that the major truncated form of CdiA lacks the C-terminal FHA-2, pretoxin and CdiA-CT domains (**Figs. 1A**), but is still presented on the cell surface as an adhesin (2, 16, 17, 62). Prior to co-culture, we treated CdiA-StbD expressing inhibitors with chloramphenicol to prevent further protein synthesis, then introduced chloramphenicol-resistant *E. coli ompF^+^ gyrB-VSV* target bacteria to initiate toxin delivery. Processed CdiA-StbD lacking the toxin region (ΔCT) is detected almost immediately after exposure to target cells (**Fig. 8B**, lanes 2-6). Concomitantly, GyrB-VSV in the target cell population is cleaved by the toxin, with nearly complete conversion within 15 min of co-culture (**Fig. 8B**, lanes 2 & 3). Gly157Ser/Gly158Ser substitutions in the linker region do not interfere with ΔCT processing, but the rate of GyrB-VSV cleavage in target bacteria is much slower (**Fig. 8B**, lanes 8-12). These results suggest that the Gly157Ser/Gly158Ser mutations have no effect on pretoxin-mediated auto-processing, but subsequent translocation to the cytosol appears to be defective. Substitution of the catalytic Cys183 residue also has no effect on ΔCT processing, but the inactivated toxin does not cleave GyrB-VSV (**Fig. 8C**, lanes 1-6). Somewhat unexpectedly, we detected a new processed form of CdiA-StbD carrying the Cys187Ser substitution. This product migrates between full-length and ΔCT forms of CdiA-StbD in immunoblots (**Fig. 8C**, lanes 7 & 8), consistent with cleavage after Gly158 in the linker between entry and C39 peptidase domains. Given that this processing occurs prior to target-cell recognition, the Cys187Ser mutation likely interferes with redox regulation, resulting in premature release of the peptidase into the inhibitor-cell periplasm. This futile process could contribute to the reduced competitive fitness of these inhibitors (**Fig. 8A**) and the incomplete cleavage of GyrB-VSV in target cells (**Fig. 8C**, lanes 8-12).

## Discussion

StbD appears to be an inactive fusion protein generated from recombination between the *fim* and *cdi* gene clusters of *C. rodentium* DBS100. Although StbD does not appear to function in inter-bacterial competition, there are many other CdiA proteins from enterobacterial and *Pseudomonas* species that carry homologous peptidase toxin domains (**Fig. S3**) (21). In addition, many *E. coli* and *Salmonella* isolates use T6SSs to deploy related C39 peptidases, exemplified by the recently described Cpe1 effector from *E. coli* ATCC 11775 (63). The latter effector carries an N-terminal PAAR-like domain, which enables the peptidase to be loaded on VgrG for export (**Fig. S4**) (63). C39 family peptidases were first identified as N-terminal extensions on ABC transporters that export lantibiotics, microcins and quorum-sensing peptides (54, 64, 65). These export proteins are referred to as ABC-transporter maturation secretion (AMS) or peptidase-containing ATP-binding transporters (PCAT) in the literature. The cytosolic C39 domains of AMS/PCATs face toward the export channel where they are positioned to cleave signal sequences after Gly-Gly/Ala/Ser motifs (64). Given that the recognition motif is relatively simple, C39 peptidases could conceivably kill bacteria through indiscriminate cleavage after diglycine sequences in many different proteins. However, the StbD-CT toxin specifically cleaves between conserved Gly residues in the ATPase domains of GyrB and ParE topoisomerase subunits. These Gly residues make direct contact with the β- and γ-phosphates of ATP (60, 66), suggesting that the toxin disrupts ATPase activity. DNA gyrase uses the free energy of ATP hydrolysis to introduce negative supercoils into the genome, whereas topoisomerase IV promotes chromosome segregation by decatenating topologically linked DNA after replication (67). *In vitro* peptidase assays suggest that GyrB may be a preferred substrate, but both type II topoisomerases are cleaved rapidly in target bacteria after toxin delivery. GyrB/ParE cleavage is coincident with decreased DNA super-helicity, and intoxicated cells become filamentous with unsegregated nucleoids, indicating that both of the essential type II topoisomerases are inactivated by StbD-CT.

Type II topoisomerases have long been known to be targets for antibiotics and antibacterial toxins. First described during the golden age of antibiotic discovery, novobiocin is an aminocoumarin that acts as a competitive inhibitor of ATP binding to GyrB/ParE (55, 68). Subsequently, quinolones were found to inhibit type II topoisomerases by blocking DNA re-ligation by the GyrA/ParC subunits (56). DNA gyrase is also poisoned by microcin B17, which is a small oxazole-thiazole modified peptide released by various enterobacterial species (69). The activity of microcin B17 is incompletely understood, but available data suggest that the peptide binds to GyrB and slows the passage of intact T-segment DNA across the cleaved G-segment (69). DNA gyrase is also inactivated by small type II protein toxins that are commonly encoded on low-copy plasmids. CcdB is a type II toxin-antitoxin (TA) encoded by the *E. coli* F plasmid that functions to stabilize the episome in bacterial populations through post-segregational killing of daughter cells that fail to inherit the plasmid (70). After Lon-mediated degradation of the CcdA antitoxin, CcdB binds to GyrA and blocks its duty cycle prior to DNA re-ligation, similar to the effect of quinolone treatment (71). More recently, another type II toxin, FicT, was shown to inactivate type II topoisomerases through enzymatic adenylylation of residue Tyr109 in the ATPase domain of GyrB (72). This conserved residue packs against the nucleobase of adenine nucleotide, while the amide of Gly102 interacts with the 2’ and 3’ hydroxyl groups of ribose (59, 61). Thus, peptidase cleavage of bacterial type II topoisomerases represents yet another strategy to inactivate these critical enzymes.

The StbD-CT•StbI binding interface is somewhat unusual compared to other characterized toxin•immunity protein complexes. Instead of physically blocking the peptidase active site, StbI binds primarily to the opposite face of the C39 domain leaving the catalytic center exposed. Immunity protein interactions with “exosites” on cognate toxins were first observed with DNase colicins E7 and E9 (73–75), and have also been reported for some CdiA-CT DNases (37, 76). This phenomenon underscores the exquisite substrate specificity of the peptidase, because the unoccluded active site does not permit adventitious cleavage of other proteins. Substrate specificity relies not only on the conserved HAGGK motif surrounding the scissile peptide, but also upon interactions with the flexible N-terminal arm of the ATPase domain. This latter segment of GyrB wraps around the peptidase, interacting with the same surface as the StbI immunity protein. Most of the substrate-binding interactions entail main-chain contacts, which may account for the toxin’s ability to recognize both GyrB and ParE. The diversification of C39 peptidase toxins also has implications for substrate specificity (**Fig. S3**). For example, the Cpe1 and StbD peptidase domains diverge at several positions that interact directly with GyrB (**Fig. S4**). There is no available structure of Cpe1 bound to GyrB, but AlphaFold 3 modeling suggests that it utilizes a similar, yet distinct, set of polar interactions from that observed with StbD-CT (**Fig. S4**) (46, 63). Although the precise bonding networks may differ between C39 peptidases, it is clear that these toxins can only cleave GyrB/ParE in the nucleotide-free state. The HAGGK motifs of GyrB/ParE make direct contact with ATP, rendering the scissile peptide bond inaccessible to the toxin. Moreover, the dimerization of ATPase domains upon binding nucleotide appears to preclude any interaction with the peptidase domain.

Finally, we find that C39 mediated auto-processing is critical for target-cell intoxication. Previous work has shown that other CdiA-CT toxins can be held within the inhibitor-cell periplasm for several hours while remaining competent for immediate delivery into target bacteria (31). Such extended dwell times could be problematic because auto-processing would release the peptidase domain before targets are encountered. Given that C39 auto-processing requires reducing agents *in vitro*, premature toxin release is likely suppressed by the oxidizing conditions that prevail in the periplasm. Once transferred into the target-cell periplasm, the StbD-CT is liberated by pretoxin-mediated cleavage after the VENN sequence, and the released toxin then searches for YajC for cell entry (**Fig. 1B**, steps 4 & 5). Given that Gly154Ser/Gly158Ser substitutions interfere with CDI-mediated intoxication, we propose that C39 auto-processing serves to release the peptidase domain upon entering the reducing environment of the cytoplasm. According to this model, the StbD-CT fragment is translocated across the cytoplasmic membrane, but remains tethered to YajC through interactions with the entry domain (**Fig. 1B**, step 5). Redox regulated auto-processing would allow the peptidase to penetrate into the cell interior where it could more effectively inactivate nucleoid associated topoisomerases. We note that the Cpe1 effector is probably subject to similar constraints because it shares the same auto-processing site as StbD-CT (**Fig. S3**). Moreover, Cpe1 contains a domain that is homologous to the StbD-CT entry domain (**Fig. S4**), suggesting that it also hijacks YajC to enter the cytosol. To our knowledge, this is the first example of a T6SS-delivered effector that utilizes a CDI toxin cell-entry pathway.

## Experimental Procedures

### Bacterial strains and growth

Bacterial strains used in this study are outlined in **Table S3**. Bacteria were cultured at 37 °C in lysogeny broth (LB) or on LB agar. Where appropriate, media were supplemented with antibiotics at the following concentrations: ampicillin (Amp) 150 µg mL^-1^, kanamycin (Kan) 50 µg mL^-1^, chloramphenicol (Cm) 33 µg mL^-1^, tetracycline (Tet) 15 µg mL^-1^ and trimethoprim (Tp) 100 µg mL^-1^. The *ΔompC::kan*, Δ*ompF::kan* and Δ*yajC::kan* deletion alleles were obtained from the Keio collection (77) and transferred into *E. coli* MG1655 Δ*wzb* (78) cells by bacteriophage P1 mediated transduction. The *pir-116* allele was transduced from *E. coli* MFD (79) into *E. coli* MC1061 using the linked Tp^R^ marker to generate strain CH8802. Where indicated, kanamycin resistance cassettes were removed by FLP recombinase expression from plasmid pCP20 (80).

### Plasmid and strain constructions

Plasmids and oligonucleotide primers are presented in **Tables S4** & **S5**, respectively. *C. rodentium* DBS100 deletion strains were generated using plasmid pCH371 (81), which carries the counter-selectable *pheS(A294G)* allele. Homology fragments flanking the *stbD-CT/stbI* coding sequence were amplified with primers **CH5527/CH5528** and **CH5529/CH5530**, followed by sequential ligation to pKAN (82) to generate plasmid pCH1863. The SacI-KpnI fragment containing the *kan* resistance cassette was subcloned from pCH1863 into pCH371 to generate plasmid pCH5918. To delete the DBS100 *cdiCAI* gene cluster, plasmid pCH1161 was generated in the same manner using primers **CH5608/CH5609** and **CH5610/CH5611** to amplify homology regions. Plasmids pCH5918 and pCH1161 were introduced into DBS100 by conjugation and exconjugants were selected on solid media supplemented with Cm and Kan. Counter-selection was conducted in M9 minimal medium supplemented with 0.4% D-glucose, 2 mM D/L-chlorophenylalanine and 50 µg/mL kanamycin. Strains CH400 (DBS100 Δ*cdiCAI::kan*) and CH1104 (DBS100 Δ*stbD-CT/stbI::kan*) were isolated as Kan^R^ Cm^S^ clones that had lost *pheS(A294G)* through homologous recombination. The rhamnose-inducible *rhaBAD* promoter was introduced upstream of the DBS100 *stbABCDI* locus using plasmid pCH501, which was generated from pRE118 (83) in two steps. The *sacB* gene of pRE118 was first removed by EcoRI digestion and religation to produce pCH461. The small SacI-SphI fragment from pSCrhaB2 (84) was then ligated to pCH461 to generate pCH501. DBS100 *stbA* was amplified with primers **CH5431**/**CH5432** and ligated to pCH501 using NcoI-SphI restriction sites to generate pCH5944. Plasmid pCH5944 was introduced into DBS100 by conjugation to produce strain CH5947. A similar procedure was used to generate a rhamnose-inducible *cdiBÁ::stbD* fusion. The *cdiBÁ* sequence was amplified from plasmid pCH13604 (22) with primers **CH5562**/**CH5561** and ligated to pCH501 via NcoI-SalI restriction sites to generate pCH3779. A fragment of *stbD* encoding residues Ile585 - Ser388 was amplified with primers **CH5560**/**CH5559** and ligated to pCH3779 via SalI-XbaI restriction sites to produce pCH3780. Plasmid pCH3780 was introduced into DBS100 by conjugation to generate strain CH4201.

The rhamnose-inducible *stbABCDI* and *cdiBÁ-stbDI* gene clusters were isolated by rescue cloning. Genomic DNA from strains CH5947 and CH4201 was digested with SphI at 37 °C for 5 h, then incubated at 75 °C for 10 min to inactivate the restriction enzyme. The digests were treated with phage T4 DNA ligase and 1 mM ATP overnight at 16 °C, and the reactions transformed into CH8802 to recover plasmids pCH946 and pCH4202. The *stbD-CT/stbI* module was also sequentially amplified with primers **CH5335/CH5338** and **CH4838/CH5338** and the final product ligated to NheI-XhoI digested pCH1285 (21) for fusion to the *E. coli* EC93 *cdiA* gene (pCH1826). Missense mutations were introduced into *stbD* using megaprimer PCR. Plasmid pCH4202 was amplified with primer **CH5899** in conjunction with **CH5425**, **CH5423**, **CH5336** and **CH5458**. The products were used as megaprimers in second reactions with primer **CH5952** and the final products ligated to KpnI-XbaI digested pCH4202 to produce plasmids pCH8908 (GG157-158SS), pCH8909 (Gly158Ser), pCH8910 (Cys183Ala) and pCH8911(Cys187Ser). Similarly, pCH4202 was amplified with **CH5952** in conjunction with primers **CH5381** and **CH5402** to generate megaprimers for final PCRs with **CH5890** to generate pCH8912 (His258Ala) and pCH8913 (Asp275Ala), respectively. The *stbD-CT/stbI* coding sequences were amplified from the latter constructs with primers **CH4346/CH4347**, and the products were ligated to NcoI/SpeI digested pCH6505 (8) to generate plasmids pCH13983 (wild-type), pCH6007 (GG157- 158SS), pCH6009 (Gly158Ser), pCH998 (Cys183Ala), pCH5997 (Cys187Ser), pCH9108 (His258Ala), and pCH9109 (Asp275Ala), which produce StbD-CT•StbI-His_6_ complexes for purification. In addition, overexpression constructs that produce StbD-CT and C39 peptidase domains with N-terminal His_6_ tags were generated with plasmid pMCSG63 (42). Wild-type *stbD-CT/stbI* was amplified with **CH5356/CH5338** and the Cys183Ala allele with **CH5356/CH5357** for ligation to KpnI-XhoI digested pMCSG63 to generate pCH1762 and pCH1764, respectively. Plasmid pCH998 was amplified with primer pairs **CH5765**/**CH5338** and **CH5765/CH5357** and the products ligated to KpnI-XhoI digested pMCSG63 to generate pCH8858 and pCH9103, respectively. The *stbI* immunity gene was amplified with primers **CH5337/CH4347** and ligated to KpnI-SpeI digested pCH8001 (85) to generate plasmid pCH9110 for the purification of StbI-His_6_. The immunity gene was also amplified with **CH5337/CH5338** for ligation to KpnI-XhoI digested pCH405Δ (17) to produce plasmid pCH9102 for immunity protection experiments.

The *ompF*^MG1655^ and *ompF*^DBS100^ genes were amplified with primers **CH2729/CH2734** and the products ligated to KpnI-XhoI digested pTrc99Cm (86) to generate plasmids pCH8210 and pCH8212, respectively. The Gly141Ala, Gly141Arg and ΔGly142-Ser147 mutations were introduced into pCH8210 by megaprimer PCR using primers **CH6643**, **CH6644** and **CH6645** in conjunction with CH2734. The resulting products were used as forward primers with **CH2729** to generate plasmids pCH8217 (Gly141Ala), pCH8218 (Gly141Arg) and pCH8219 (ΔGly142-Ser147). The *ompC*^MG1655^ coding sequence was amplified with **CH2387/CH2388** and ligated to KpnI-XhoI digested pTrc99Cm to generate pCH8211. The *yajC* gene was amplified with **CH4434/CH4435** and ligated to KpnI-XhoI digested pTrc99a to generate pCH8510.

Topoisomerase subunits were tagged with VSV-G epitopes using a derivative of plasmid pCH371. The EcoRI-KpnI fragment encoding *vgrG2-VSV* was subcloned from pCH14397 (87) into pCH371 to generate integrative plasmid pCH5955. Homology fragments were amplified for *gyrA* (**CH5694**/**CH5696**), *gyrB* (**CH5698**/**CH5700**), *parC* (**CH5767**/**CH5768**) and *parE* (**CH5702**/**CH5766**), and the products ligated to EcoRI-SpeI digested pCH5955 to generate plasmids pCH7987, pCH7988, pCH7990 and pCH7991, respectively. These constructs were introduced into *E. coli* CH7367 by conjugation to generate strains CH7993 (*gyrA-VSV*), CH7994 (*gyrB-VSV*), CH7992 (*parC-VSV*) and CH7995 (*parE-VSV*). Individual topoisomerase subunit genes were amplified with **CH5695**/**CH5697** (*gyrA*), **CH5699**/**CH5701** (*gyrB*) and **CH5715**/**CH5716** (*parE*). The resulting products were digested with NheI-XhoI and ligated to pET21b to generate plasmids pCH8187, pCH8189 and pCH8191, respectively. Coding sequences for the N-terminal ATPase domains of GyrB and ParE were amplified with primers **CH5699**/**CH5714** and **CH5715**/**CH5828** and ligated to NheI-XhoI digested pET21b to generate plasmids pCH8193 and pCH8513, respectively. Site-directed mutations were introduced into the GyrB ATPase domain using oligonucleotides **CH5824** (Gly101Ser), **CH5825** (Gly102Ser), **CH5748** (Gly101Ser/Gly102Ser) and **CH5749** (Gly113Ser/Gly114Ser) in megaprimer PCRs with **CH5699**/**CH5714** to generate plasmids pCH8511, pCH8512, pCH5358 and pCH5359, respectively. The N-terminal arm of GyrB was deleted using primers **CH6082**/**CH5714** followed by ligation to KpnI-XhoI digested pET21K to produce plasmid pCH4032.

### Competition co-cultures

Inhibitor strains were grown to mid-log phase at 37 °C in LB medium supplemented with Kan and 0.4% L-rhamnose. *E. coli* and *C. rodentium* target bacteria were cultured to mid-log phase at 37 °C in LB media supplemented with either Amp or Cm. Inhibitor and target bacteria were adjusted to an optical density at 600 nm (OD_600_) of 3.0 in LB media supplemented with 0.4% L-rhamnose, mixed at a 1:1 ratio and plated onto LB agar for 3 h at 37 °C. Bacteria were harvested with sterile swabs into 1.0 mL of M9 salts, and serial dilutions were plated onto antibiotic selective media to enumerate viable cells as colony forming units for both populations. Competitive indices were calculated as the ratio of inhibitors to targets at 3 h divided by the initial ratio. Presented data are the averages ± standard error (SEM) for three independent experiments.

### Mutagenesis and CDI^R^ selections

*mariner* transposon mutant pools were generated as described previously (23). MFD *pir*^+^ donor cells carrying plasmid pSC189 were grown in LB medium supplemented with Amp and 30 µM diaminopimelic acid, then mixed with *E. coli* CH7367 recipients on LB agar at 37 °C for 4 h (79, 88). Cell mixtures were harvested using a sterile polyester swab and plated onto Kan-supplemented LB agar to select for transposon mutants. Six independent mutant pools were each harvested into 1 mL of 1X M9 salts for subsequent selection of StbD-CT resistant mutants. Inhibitors carrying plasmid pCH1826 were co-cultured at a 1:2 ratio with each *mariner* mutant pool in 25 mL of LB media for 3 h at 37 °C. Surviving target bacteria were isolated on LB agar supplemented with Kan for further selection by iterative competition co-culture. After three rounds selection, individual clones were tested for resistance. Transposon insertion sites were identified by rescue cloning. Approximately 1 µg of genomic DNA from each resistant mutant was digested with NspI at 37 °C overnight, followed by inactivation of the restriction enzyme at 65 °C for 25 min. Heat-inactivated reactions were supplemented with 1 mM ATP and T4 DNA ligase for overnight incubation at 16 °C. Plasmids were recovered from Kan-resistant *E. coli* DH5ɑ *pir^+^* transformants, and *mariner* insertion sites identified by DNA sequence analysis using primer CH2260.

### Protein purification

All proteins for *in vitro* biochemical analyses were produced in *E. coli* CH2016 (89). Cells were grown to OD_600_ ∼ 0.6 in Amp-supplemented LB media at 37 °C, and protein production was induced for 1 h with 1.5 mM isopropyl β-D-1-thiogalactopyranoside (IPTG). Cultures were harvested by centrifugation at 5,000 rpm at 4 °C, and the cells frozen at −80 °C. Frozen cell pellets were resuspended in 20 mM Tris-HCl (pH 7.5), 150 mM NaCl, 25 mM imidazole supplemented with 1 mg/mL lysozyme for 30 min. Suspensions were sonicated to break cells and debris was removed by centrifugation at 15,000 rpm for 15 min. Ni^2+^-nitrilotriacetic acid (Ni^2+^-NTA) agarose resin was added to cleared supernatants to bind His_6_-tagged proteins for 1 h at 4 °C. All resins were washed with 10 volumes of 20 mM Tris-HCl (pH 7.5), 150 mM NaCl, 25 mM imidazole before elution 250 mM imidazole. For the purification of untagged StbD-CT toxins, the His_6_-tagged immunity protein was removed from the complex under denaturing conditions with 6 M guanidine-HCl, 20 mM Tris-HCl (pH 7.5) prior to elution of the toxin with 250 mM imidazole. Purified proteins were dialyzed against 20 mM Tris-HCl (pH 7.5), 150 mM NaCl for storage. Protein concentrations were determined using extinction coefficients at 280 nm of 61,210 M^-1^ cm^-1^ (StbD-CT); 43,970 M^-1^ cm^-1^ (C39 domain); 14,440 M^-1^ cm^-1^ (StbI-His_6_); 52,260 M^-1^ cm^-1^ (GyrA); 71,740 M^-1^ cm^-1^ (GyrB); 66,350 M^-1^ cm^-1^ (ParE); 14,400 M^-1^ cm^-1^ (GyrB ATPase domain); 12,950 M^-1^ cm^-1^ (ΔS2-K14 ATPase domain).

### C39 auto-processing and *in vitro* peptidase assays

StbD-CT (1 µM) auto-processing was performed for 30 min at 37 °C in 20 mM Tris-HCl (pH 7.5), 150 mM NaCl after addition of dithiothreitol (DTT) to 1 mM final concentration. Equimolar StbI-His_6_ was included where indicated. Reactions were quenched with SDS gel loading buffer and analyzed by SDS-PAGE. In separate reactions, the N-terminal auto-processing fragment was isolated by Ni^2+^-affinity chromatography, then purified by reverse phase high performance liquid chromatography on a Vydac C4 column (15 x 300 mm) in 0.06% trifluoroacetic acid. The column was developed with a linear gradient of 0.052% trifluoroacetic acid in 80% acetonitrile over 60 min. The isolated peptide was dried and redissolved in formic acid for electrospray ionization-mass spectrometry. Topoisomerase cleavage assays were conducted under the same conditions in 20 mM Tris-HCl (pH 7.5), 150 mM NaCl, 1 mM DTT.

For GyrB-C39 binding experiments, the N-terminal His_6_-tag was removed from the inactive C39(Cys183Ala) domain using tobacco etch virus (TEV) protease. His_6_-tagged GyrB ATPase domains (5 µM) were mixed with C39(Cys183Ala) (5 µM) in 600 µL of 20 mM Tris-HCl (pH 7.5), 150 mM NaCl and incubated at 4 °C for 1 h. Aliquots (15 µL) were removed as input samples before addition of Ni^2+^-NTA resin (50 µL) for 30 min at 4 °C. After centrifugation, the supernatant was collected as the unbound fraction. Resins were batch washed with 20 mM Tris-HCl (pH 7.5), 150 mM NaCl and bound proteins eluted with 20 mM Tris-HCl (pH 7.5), 150 mM NaCl, 250 mM imidazole. Input, unbound and eluted samples were analyzed by SDS-PAGE.

### Crystallography

Inactive StbD-CT(C183A) was produced from plasmid pCH1764 in *E. coli* BL21-Gold(DE3) cultured in LB supplemented with Amp at 50 µg/mL. At OD_600_ ∼ 1.0, expression was induced with 0.5 mM IPTG and the culture incubated overnight at 18 °C. Cells were collected by centrifugation for 15 min at 5,000 rpm, and the cell pellet frozen at −20 °C. Frozen cells were resuspended in 50 mM Tris-HCl (pH 7.5), 300 mM NaCl, 10% glycerol (Buffer A) supplemented with 1 mM phenylmethylsulfonyl fluoride (PMSF) and 100 µg/mL lysozyme for sonication. Debris was removed by centrifugation at 14,000 rpm for 1 h, and the supernatant applied to a HisTrap FF column (GE Healthcare) in Buffer A at 4 °C. Protein was eluted with a linear gradient of 0 to 500 mM imidazole in Buffer A. Fractions containing StbD-CT(C183A) were identified by SDS-PAGE. Pooled fractions were dialyzed against 20 mM Tris-HCl (pH 7.5), 150 mM NaCl at 4 °C and concentrated to 30 mg/mL for crystallization trials.

After seven months at 4 °C, protein crystals grew in JCSG 35 [0.1 M sodium acetate (pH 4.6), 2 M ammonium sulfate]. The crystal condition was optimized to 0.1 M sodium acetate (pH 4.0), 2 M ammonium sulfate with 1:1 protein to well drops. After 3 months at 4 °C, crystals were cryo-cooled in mother liquor supplemented with 15% glycerol and sent to SSRL 12-2 for remote data collection. Both native (E = 12,398 eV) and S-SAD (E = 7,000 eV) data were collected (**Table 1**). The native 1.9 Å dataset was not sufficient to determine the structure of StbD-CT(C183A) by molecular replacement (MR) with homologous structures or homology-based models. S-SAD data were collected from three separate crystals, one of which was the source of the native dataset. The data were processed in XDS (90), and the best frames were scaled together in XSCALE (17,540 frames in total), before processing in AIMLESS (91). The resulting data diffracted to 2 Å and had anomalous signal to 3.5 Å. Although MR was not successful in the absence of S-SAD data, a phaser MR search based on the AlphaFold model (46) was successful with the incorporation of S-SAD data in AutoSol (92, 93). The resulting model was then improved in AutoBuild (94), before subsequent rounds of manual refinement in Coot and computational refinement in phenix.refine (95, 96). The refined MR-SAD model was used for MR in phaser with the 1.9 Å native dataset described above (92), and refined in phenix.refine and Coot (95, 96). Refinement in phenix.refine utilized Translation/Libration/Screw (TLS) rotation parameters where each subunit was treated as two TLS groups: chain A and C (residues 13-160, 162-318) and chain B and D (residues 13-84, 85-318). The structure was refined to a final R_work_/R_free_ of 16.2/18.6 (**Table 1**).

The C39(C183A)•StbI complex was produced from plasmid pCH8858 in *E. coli* BL21-Gold(DE3) cultured in LB supplemented with Amp at 50 µg/mL. At OD_600_ ∼ 0.9, expression was induced with 0.5 mM IPTG and the culture incubated overnight at 18 °C. Cells were collected by centrifugation for 15 min at 5,000 rpm and the cell pellet frozen at −20 °C. Cell were broken by sonication in 20 mM sodium phosphate (pH 7.4), 150 mM NaCl, 10 % glycerol (Buffer B) supplemented with PMSF and lysozyme. Debris was removed by centrifugation at 14,000 rpm for 1 h, and the supernatant applied to a HisTrap FF column (GE Healthcare) in Buffer B at 4 °C. Protein was eluted with a linear gradient of 0 to 500 mM imidazole in Buffer B. Fractions containing the complex were dialyzed against 50 mM Tris-HCl (pH 7.5), 300 mM sodium chloride, 1 mM DTT in the presence of TEV protease to remove the N-terminal His_6_- tag from the C39 peptidase domain. The treated complex passed over Ni^2+^-NTA agarose to remove His_6_-tagged TEV protease, and the flow-through was concentrated to 30 mg/mL for crystallization trials at 4 °C. The C39(C183A)•StbI complex was screened for crystallization at 4 °C in sparse matrix screens. After two weeks, crystals were detected in Index 73 [0.1 M Tris-HCl (pH 8.5), 0.2 M sodium chloride, 25 % PEG 3350]. The crystals were cryo-cooled in mother liquor supplemented with 15% glycerol directly from the screen condition. A native dataset was collected at ALS 5.0.2 with resolution to 2.5 Å (**Table 1**). The dataset was solved by MR in phaser (92) using the C39 domain from the StbD-CT(C183A) structure (PDB:8SSK) and an AlphaFold model of StbI (46). The structure was refined with rounds of manual refinement in Coot and computational refinement in phenix.refine to a final R_work_/R_free_ of 18.7/23.7 (**Table 1**).

The N-terminal ATPase domain of GyrB was produced from plasmid pCH8193 and the inactive C39(Cys183Ala) peptidase was produced from pCH9103 in *E. coli* BL21-Gold(DE3) cultured in terrific broth supplemented with Amp at 100 µg/mL. At OD_600_ ∼ 0.6, expression was induced with 0.5 mM IPTG for overnight incubation at 20 °C. Cells were collected by centrifugation for 15 min at 5,000 rpm and the cell pellet frozen at −20 °C. Cell were broken by sonication in Buffer B supplemented with PMSF and lysozyme. Debris was removed by centrifugation at 14,000 rpm for 1 h, and the supernatant clarified by filtration. Lysate was applied to a HisTrap column (Cytiva, Marlborough, MA) pre-equilibrated with Buffer B using a Bio-Rad Biologic FPLC instrument (Hercules, CA). Following washing with 10 column volumes of Buffer B, the column was developed with a linear gradient of 0 to 500 mM imidazole in Buffer B. Fractions containing pure GyrB ATPase domain and C39(Cys183Ala) peptidase were concentrated using an Amicon 30 kDa cut-off concentrator (MilliporeSigma, Burlington, MA) in Buffer B.

Purified GyrB ATPase domain (15 mg/mL) with inactive C39(C183A) peptidase (15 mg/mL) were combined for crystallization trials. An initial crystal hit was found after one week at room temperature in Index condition 42 [0.1M Bis-Tris (pH 5.5), 25% PEG-3350] (Hampton Research). Crystallization conditions were optimized, and the best crystals were reproduced under the same condition with a 1:1 ratio of protein complex to well solution. C39•GyrB complex crystals were cryocooled in mother liquor supplemented by 20% glycerol after three weeks. Diffraction data were collected remotely at SSRL and processed in XDS to a final resolution of 2 Å (90). A model was determined by molecular replacement in phaser (92) using the previously solved C39 structure (PDB: 8SSM) and an AlphaFold generated model for the GyrB ATPase domain (46). The resulting complex model was improved in AutoBuild (94), and then refined manually in coot and computationally in phenix.refine to a final R_work_/R_free_ of 17.5/21.5 (**Table 1**) (95, 96).

### Target cell intoxication analysis

sfYFP labeled inhibitor cells (CH2445) that deploy StbD-CT from plasmid pCH1826 were co-cultured with mKate-labeled target bacteria (CH2567) (76) that carry pCH405Δ (*stbI^-^*) or pCH9102 (*stbI^+^*) in LB media at 37 °C. Cells were collected by centrifugation, washed with 1 mL of PBS and resuspended in 100 μL of PBS supplemented with 1 μg/mL of Hoechst 33342 for 5 min. Cells were then washed with PBS and resuspended in 100 µL of PBS. Cell suspensions (5 μL) were spotted onto 1% agarose pads for visualization on an ECHO Revolve R4 microscope under 100x objective lens with immersion oil. CH8896 inhibitors that carry pCH4202 and pCH8910 were grown to mid-log phase in LB media supplemented with 0.2% L-rhamnose, then mixed at a 1:1 ratio with target bacteria that harbor plasmid pBluescript II SK(+). After 30 min at 37 °C, the co-cultures were harvested by centrifugation and plasmid DNA isolated. Plasmids were electrophoresed on a 1% agarose gel supplemented with 1 µg/mL chloroquine diphosphate at 100 V for 4 h (97). The gel was stained with 1 µg/mL ethidium bromide for 30 min, then washed three times with distilled water before imaging on a UV lightbox.

### Immunoblot analysis of co-cultures

Inhibitor cells carrying plasmid pCH4202 were grown to mid-log phase at 37 °C in LB medium supplemented with Kan and 0.2% L-rhamnose. *E. coli* target strains (CH7992, CH7993, CH7994 and CH7995) were grown to mid-log phase at 37 °C in LB media supplemented with Cm. Inhibitor cell cultures were treated with Cm (66 µg/mL) for 30 min before mixing with target bacteria at a 1:2 cell ratio in Cm-supplemented LB media. Co-cultures were incubated at 37 °C with shaking and samples collected at the indicated time points. Cell pellets were frozen at –80 °C for freeze-thaw lysis in 50 µL of 8 M urea lysis buffer [8 M urea, 20 mM Tris-HCl (pH 8.0), 150 mM NaCl]. Protein concentrations were determined by Bradford assay and equal amounts of each sample were electrophoresed on 6% polyacrylamide SDS gels run at 100 V (constant) for 1.5 h. Gels were soaked for 5 min in 25 mM Tris, 192 mM glycine (pH 8.6), 10% methanol, then electroblotted to polyvinylidene fluoride membranes using a semi-dry transfer apparatus at 17 V for 1 h. Membranes were blocked with 4% non-fat milk in PBS for 1 h at room temperature and incubated with polyclonal antibodies (1:5,000 dilution) to the N-terminal TPS domain of CdiA (22, 62) and mouse monoclonal antibody to the VSV-G epitope (Sigma-Aldrich, 1:5,000 dilution) in 0.1% not-fat milk in PBS overnight at 4 °C. Blots were incubated with IRDye^®^ 800CW-conjugated goat anti-rabbit IgG (1:25,000 dilution, LICOR) and IRDye^®^ 680LT-conjugated goat anti-mouse IgG (1:25,000 dilution, LICOR) in PBS. Immunoblots were visualized with an Azure Biosystems Sapphire FL Biomolecular Imager.

## Acknowledgments

We thank Greg Ekberg for generating plasmid pCH13984 and Jessica Mendoza for technical support. This work was supported by National Institutes of Health grants GM117373 and GM144437 (C.W.G & C.S.H.).

## References

1. Ruhe ZC, Low DA, Hayes CS. 2020. Polymorphic Toxins and Their Immunity Proteins: Diversity, Evolution, and Mechanisms of Delivery. Annu Rev Microbiol 74:497–520.

2. Aoki SK, Pamma R, Hernday AD, Bickham JE, Braaten BA, Low DA. 2005. Contact-dependent inhibition of growth in *Escherichia coli*. Science 309:1245–8.

3. Hood RD, Singh P, Hsu F, Guvener T, Carl MA, Trinidad RR, Silverman JM, Ohlson BB, Hicks KG, Plemel RL, Li M, Schwarz S, Wang WY, Merz AJ, Goodlett DR, Mougous JD. 2010. A type VI secretion system of *Pseudomonas aeruginosa* targets a toxin to bacteria. Cell Host Microbe 7:25–37.

4. MacIntyre DL, Miyata ST, Kitaoka M, Pukatzki S. 2010. The *Vibrio cholerae* type VI secretion system displays antimicrobial properties. Proc Natl Acad Sci U S A 107:19520–4.

5. Souza DP, Oka GU, Alvarez-Martinez CE, Bisson-Filho AW, Dunger G, Hobeika L, Cavalcante NS, Alegria MC, Barbosa LR, Salinas RK, Guzzo CR, Farah CS. 2015. Bacterial killing via a type IV secretion system. Nat Commun 6:6453.

6. Garcia-Bayona L, Guo MS, Laub MT. 2017. Contact-dependent killing by *Caulobacter crescentus* via cell surface-associated, glycine zipper proteins. Elife 6.

7. Vassallo CN, Cao P, Conklin A, Finkelstein H, Hayes CS, Wall D. 2017. Infectious polymorphic toxins delivered by outer membrane exchange discriminate kin in myxobacteria. Elife 6.

8. Aoki SK, Diner EJ, de Roodenbeke CT, Burgess BR, Poole SJ, Braaten BA, Jones AM, Webb JS, Hayes CS, Cotter PA, Low DA. 2010. A widespread family of polymorphic contact-dependent toxin delivery systems in bacteria. Nature 468:439–42.

9. De Gregorio E, Esposito EP, Zarrilli R, Di Nocera PP. 2018. Contact-Dependent Growth Inhibition Proteins in *Acinetobacter baylyi* ADP1. Curr Microbiol 75:1434–1440.

10. Mercy C, Ize B, Salcedo SP, de Bentzmann S, Bigot S. 2016. Functional Characterization of *Pseudomonas* Contact Dependent Growth Inhibition (CDI) Systems. PLoS One 11:e0147435.

11. Beck CM, Morse RP, Cunningham DA, Iniguez A, Low DA, Goulding CW, Hayes CS. 2014. CdiA from *Enterobacter cloacae* delivers a toxic ribosomal RNase into target bacteria. Structure 22:707–718.

12. Clantin B, Delattre AS, Rucktooa P, Saint N, Meli AC, Locht C, Jacob-Dubuisson F, Villeret V. 2007. Structure of the membrane protein FhaC: a member of the Omp85-TpsB transporter superfamily. Science 317:957–61.

13. Guerin J, Botos I, Zhang Z, Lundquist K, Gumbart JC, Buchanan SK. 2020. Structural insight into toxin secretion by contact-dependent growth inhibition transporters. Elife 9:e58100.

14. Aoki SK, Malinverni JC, Jacoby K, Thomas B, Pamma R, Trinh BN, Remers S, Webb J, Braaten BA, Silhavy TJ, Low DA. 2008. Contact-dependent growth inhibition requires the essential outer membrane protein BamA (YaeT) as the receptor and the inner membrane transport protein AcrB. Mol Microbiol 70:323–40.

15. Ruhe ZC, Wallace AB, Low DA, Hayes CS. 2013. Receptor polymorphism restricts contact-dependent growth inhibition to members of the same species. MBio 4:e00480–13.

16. Ruhe ZC, Nguyen JY, Xiong J, Koskiniemi S, Beck CM, Perkins BR, Low DA, Hayes CS. 2017. CdiA effectors use modular receptor-binding domains to recognize target bacteria. MBio 8:e00290–17.

17. Halvorsen TM, Garza-Sánchez F, Ruhe ZC, Bartelli NL, Chan NA, Nguyen JY, Low DA, Hayes CS. 2021. Lipidation of Class IV CdiA Effector Proteins Promotes Target Cell Recognition during Contact-Dependent Growth Inhibition. mBio 12:e0253021.

18. Beck CM, Willett JL, Cunningham DA, Kim JJ, Low DA, Hayes CS. 2016. CdiA Effectors from Uropathogenic *Escherichia coli* Use Heterotrimeric Osmoporins as Receptors to Recognize Target Bacteria. PLoS Pathog 12:e1005925.

19. Nikolakakis K, Amber S, Wilbur JS, Diner EJ, Aoki SK, Poole SJ, Tuanyok A, Keim PS, Peacock S, Hayes CS, Low DA. 2012. The toxin/immunity network of *Burkholderia pseudomallei* contact-dependent growth inhibition (CDI) systems. Mol Microbiol 84:516–29.

20. Aoki SK, Webb JS, Braaten BA, Low DA. 2009. Contact-dependent growth inhibition causes reversible metabolic downregulation in *Escherichia coli*. J Bacteriol 191:1777–86.

21. Halvorsen TM, Schroeder KA, Jones AM, Hammarlöf D, Low DA, Koskiniemi S, Hayes CS. 2024. Contact-dependent growth inhibition (CDI) systems deploy a large family of polymorphic ionophoric toxins for inter-bacterial competition. PLoS Genet 20:e1011494.

22. Ruhe ZC, Subramanian P, Song K, Nguyen JY, Stevens TA, Low DA, Jensen GJ, Hayes CS. 2018. Programmed Secretion Arrest and Receptor-Triggered Toxin Export during Antibacterial Contact-Dependent Growth Inhibition. Cell 175:921–933 e14.

23. Willett JL, Gucinski GC, Fatherree JP, Low DA, Hayes CS. 2015. Contact-dependent growth inhibition toxins exploit multiple independent cell-entry pathways. Proc Natl Acad Sci U S A 112:11341–6.

24. Allen JP, Hauser AR. 2019. Diversity of *Pseudomonas aeruginosa* contact-dependent growth inhibition systems. J Bacteriol doi:10.1128/JB.00776-18.

25. Guerin J, Bigot S, Schneider R, Buchanan SK, Jacob-Dubuisson F. 2017. Two-Partner Secretion: Combining Efficiency and Simplicity in the Secretion of Large Proteins for Bacteria-Host and Bacteria-Bacteria Interactions. Front Cell Infect Microbiol 7:148.

26. Baud C, Guerin J, Petit E, Lesne E, Dupre E, Locht C, Jacob-Dubuisson F. 2014. Translocation path of a substrate protein through its Omp85 transporter. Nat Commun 5:5271.

27. Guerin J, Saint N, Baud C, Meli AC, Etienne E, Locht C, Vezin H, Jacob-Dubuisson F. 2015. Dynamic interplay of membrane-proximal POTRA domain and conserved loop L6 in Omp85 transporter FhaC. Mol Microbiol 98:490–501.

28. Kajava AV, Cheng N, Cleaver R, Kessel M, Simon MN, Willery E, Jacob-Dubuisson F, Locht C, Steven AC. 2001. Beta-helix model for the filamentous haemagglutinin adhesin of *Bordetella pertussis* and related bacterial secretory proteins. Mol Microbiol 42:279–92.

29. Mirdita M, Schütze K, Moriwaki Y, Heo L, Ovchinnikov S, Steinegger M. 2022. ColabFold: making protein folding accessible to all. Nat Methods 19:679–682.

30. Tiu AKY, Conroy GC, Bobst CE, Hagan CL. 2024. Autoproteolytic mechanism of CdiA toxin release reconstituted in vitro. J Bacteriol 206:e0024924.

31. Ruhe ZC, Nguyen JY, Beck CM, Low DA, Hayes CS. 2014. The proton-motive force is required for translocation of CDI toxins across the inner membrane of target bacteria. Mol Microbiol 94:466–81.

32. Bartelli NL, Sun S, Gucinski GC, Zhou H, Song K, Hayes CS, Dahlquist FW. 2019. The Cytoplasm-Entry Domain of Antibacterial CdiA Is a Dynamic alpha-Helical Bundle with Disulfide-Dependent Structural Features. J Mol Biol 431:3203–3216.

33. Bartelli NL, Passanisi VJ, Michalska K, Song K, Nhan DQ, Zhou H, Cuthbert BJ, Stols LM, Eschenfeldt WH, Wilson NG, Basra JS, Cortes R, Noorsher Z, Gabraiel Y, Poonen-Honig I, Seacord EC, Goulding CW, Low DA, Joachimiak A, Dahlquist FW, Hayes CS. 2022. Proteolytic processing induces a conformational switch required for antibacterial toxin delivery. Nat Commun 13:5078.

34. Koskiniemi S, Garza-Sanchez F, Edman N, Chaudhuri S, Poole SJ, Manoil C, Hayes CS, Low DA. 2015. Genetic analysis of the CDI pathway from *Burkholderia pseudomallei* 1026b. PLoS One 10:e0120265.

35. Myers-Morales T, Sim MMS, DuCote TJ, Garcia EC. 2021. *Burkholderia multivorans* requires species-specific GltJK for entry of a contact-dependent growth inhibition system protein. Mol Microbiol 116:957–973.

36. Allen JP, Ozer EA, Minasov G, Shuvalova L, Kiryukhina O, Satchell KJF, Hauser AR. 2020. A comparative genomics approach identifies contact-dependent growth inhibition as a virulence determinant. Proc Natl Acad Sci U S A 117:6811–6821.

37. Morse RP, Nikolakakis KC, Willett JL, Gerrick E, Low DA, Hayes CS, Goulding CW. 2012. Structural basis of toxicity and immunity in contact-dependent growth inhibition (CDI) systems. Proc Natl Acad Sci U S A 109:21480–21485.

38. Johnson PM, Gucinski GC, Garza-Sanchez F, Wong T, Hung LW, Hayes CS, Goulding CW. 2016. Functional Diversity of Cytotoxic tRNase/Immunity Protein Complexes from *Burkholderia pseudomallei*. J Biol Chem 291:19387–400.

39. Batot G, Michalska K, Ekberg G, Irimpan EM, Joachimiak G, Jedrzejczak R, Babnigg G, Hayes CS, Joachimiak A, Goulding CW. 2017. The CDI toxin of *Yersinia kristensenii* is a novel bacterial member of the RNase A superfamily. Nucleic Acids Res 45:5013–5025.

40. Michalska K, Gucinski GC, Garza-Sanchez F, Johnson PM, Stols LM, Eschenfeldt WH, Babnigg G, Low DA, Goulding CW, Joachimiak A, Hayes CS. 2017. Structure of a novel antibacterial toxin that exploits elongation factor Tu to cleave specific transfer RNAs. Nucleic Acids Res 45:10306–10320.

41. Michalska K, Quan Nhan D, Willett JLE, Stols LM, Eschenfeldt WH, Jones AM, Nguyen JY, Koskiniemi S, Low DA, Goulding CW, Joachimiak A, Hayes CS. 2018. Functional plasticity of antibacterial EndoU toxins. Mol Microbiol 109:509–527.

42. Gucinski GC, Michalska K, Garza-Sanchez F, Eschenfeldt WH, Stols L, Nguyen JY, Goulding CW, Joachimiak A, Hayes CS. 2019. Convergent Evolution of the Barnase/EndoU/Colicin/RelE (BECR) Fold in Antibacterial tRNase Toxins. Structure 27:1660–1674.

43. Zhang D, de Souza RF, Anantharaman V, Iyer LM, Aravind L. 2012. Polymorphic toxin systems: Comprehensive characterization of trafficking modes, processing, mechanisms of action, immunity and ecology using comparative genomics. Biol Direct 7:18.

44. Rice AJ, Park A, Pinkett HW. 2014. Diversity in ABC transporters: type I, II and III importers. Crit Rev Biochem Mol Biol 49:426–37.

45. Virtanen P, Wäneskog M, Koskiniemi S. 2019. Class II contact-dependent growth inhibition (CDI) systems allow for broad-range cross-species toxin delivery within the Enterobacteriaceae family. Mol Microbiol 111:1109–1125.

46. Abramson J, Adler J, Dunger J, Evans R, Green T, Pritzel A, Ronneberger O, Willmore L, Ballard AJ, Bambrick J, Bodenstein SW, Evans DA, Hung CC, O’Neill M, Reiman D, Tunyasuvunakool K, Wu Z, Žemgulytė A, Arvaniti E, Beattie C, Bertolli O, Bridgland A, Cherepanov A, Congreve M, Cowen-Rivers AI, Cowie A, Figurnov M, Fuchs FB, Gladman H, Jain R, Khan YA, Low CMR, Perlin K, Potapenko A, Savy P, Singh S, Stecula A, Thillaisundaram A, Tong C, Yakneen S, Zhong ED, Zielinski M, Žídek A, Bapst V, Kohli P, Jaderberg M, Hassabis D, Jumper JM. 2024. Accurate structure prediction of biomolecular interactions with AlphaFold 3. Nature 630:493–500.

47. Benson SA, Decloux A. 1985. Isolation and characterization of outer membrane permeability mutants in Escherichia coli K-12. J Bacteriol 161:361–7.

48. Benson SA, Occi JL, Sampson BA. 1988. Mutations that alter the pore function of the OmpF porin of Escherichia coli K12. J Mol Biol 203:961–70.

49. Housden NG, Hopper JT, Lukoyanova N, Rodriguez-Larrea D, Wojdyla JA, Klein A, Kaminska R, Bayley H, Saibil HR, Robinson CV, Kleanthous C. 2013. Intrinsically disordered protein threads through the bacterial outer-membrane porin OmpF. Science 340:1570–4.

50. Francis MR, Webby MN, Housden NG, Kaminska R, Elliston E, Chinthammit B, Lukoyanova N, Kleanthous C. 2021. Porin threading drives receptor disengagement and establishes active colicin transport through Escherichia coli OmpF. Embo j 40:e108610.

51. Pogliano KJ, Beckwith J. 1994. Genetic and molecular characterization of the Escherichia coli secD operon and its products. J Bacteriol 176:804–14.

52. Duong F, Wickner W. 1997. Distinct catalytic roles of the SecYE, SecG and SecDFyajC subunits of preprotein translocase holoenzyme. Embo j 16:2756–68.

53. Ishii S, Yano T, Ebihara A, Okamoto A, Manzoku M, Hayashi H. 2010. Crystal Structure of the Peptidase Domain of *Streptococcus* ComA, a Bifunctional ATP-binding Cassette Transporter Involved in the Quorum-sensing Pathway. Journal of Biological Chemistry 285:10777–10785.

54. Bobeica SC, Dong S-H, Huo L, Mazo N, McLaughlin MI, Jiménez-Osés G, Nair SK, van der Donk WA. 2019. Insights into AMS/PCAT transporters from biochemical and structural characterization of a double Glycine motif protease. eLife 8:e42305.

55. Maxwell A. 1993. The interaction between coumarin drugs and DNA gyrase. Mol Microbiol 9:681–6.

56. Pham TDM, Ziora ZM, Blaskovich MAT. 2019. Quinolone antibiotics. Medchemcomm 10:1719–1739.

57. Bates AD, Berger JM, Maxwell A. 2011. The ancestral role of ATP hydrolysis in type II topoisomerases: prevention of DNA double-strand breaks. Nucleic Acids Res 39:6327–39.

58. Basu A, Parente AC, Bryant Z. 2016. Structural Dynamics and Mechanochemical Coupling in DNA Gyrase. J Mol Biol 428:1833–45.

59. Wigley DB, Davies GJ, Dodson EJ, Maxwell A, Dodson G. 1991. Crystal structure of an N-terminal fragment of the DNA gyrase B protein. Nature 351:624–9.

60. Vanden Broeck A, Lotz C, Ortiz J, Lamour V. 2019. Cryo-EM structure of the complete *E. coli* DNA gyrase nucleoprotein complex. Nat Commun 10:4935.

61. Brino L, Urzhumtsev A, Mousli M, Bronner C, Mitschler A, Oudet P, Moras D. 2000. Dimerization of *Escherichia coli* DNA-gyrase B provides a structural mechanism for activating the ATPase catalytic center. J Biol Chem 275:9468–75.

62. Ruhe ZC, Townsley L, Wallace AB, King A, Van der Woude MW, Low DA, Yildiz FH, Hayes CS. 2015. CdiA promotes receptor-independent intercellular adhesion. Mol Microbiol 98:175–92.

63. Song PY, Tsai CE, Chen YC, Huang YW, Chen PP, Wang TH, Hu CY, Chen PY, Ku C, Hsia KC, Ting SY. 2025. An interbacterial cysteine protease toxin inhibits cell growth by targeting type II DNA topoisomerases GyrB and ParE. PLoS Biol 23:e3003208.

64. Beis K, Rebuffat S. 2019. Multifaceted ABC transporters associated to microcin and bacteriocin export. Res Microbiol 170:399–406.

65. Håvarstein LS, Diep DB, Nes IF. 1995. A family of bacteriocin ABC transporters carry out proteolytic processing of their substrates concomitant with export. Mol Microbiol 16:229–40.

66. Lewis RJ, Singh OM, Smith CV, Skarzynski T, Maxwell A, Wonacott AJ, Wigley DB. 1996. The nature of inhibition of DNA gyrase by the coumarins and the cyclothialidines revealed by X-ray crystallography. EMBO J 15:1412–20.

67. Zechiedrich EL, Khodursky AB, Cozzarelli NR. 1997. Topoisomerase IV, not gyrase, decatenates products of site-specific recombination in *Escherichia coli*. Genes Dev 11:2580–92.

68. Bisacchi GS, Manchester JI. 2015. A New-Class Antibacterial-Almost. Lessons in Drug Discovery and Development: A Critical Analysis of More than 50 Years of Effort toward ATPase Inhibitors of DNA Gyrase and Topoisomerase IV. ACS Infect Dis 1:4–41.

69. Collin F, Maxwell A. 2019. The Microbial Toxin Microcin B17: Prospects for the Development of New Antibacterial Agents. J Mol Biol 431:3400–3426.

70. Bernard P, Couturier M. 1992. Cell killing by the F plasmid CcdB protein involves poisoning of DNA-topoisomerase II complexes. J Mol Biol 226:735–45.

71. Bahassi EM, O’Dea MH, Allali N, Messens J, Gellert M, Couturier M. 1999. Interactions of CcdB with DNA gyrase. Inactivation of GyrA, poisoning of the gyrase-DNA complex, and the antidote action of CcdA. J Biol Chem 274:10936–44.

72. Harms A, Stanger FV, Scheu PD, de Jong IG, Goepfert A, Glatter T, Gerdes K, Schirmer T, Dehio C. 2015. Adenylylation of Gyrase and Topo IV by FicT Toxins Disrupts Bacterial DNA Topology. Cell Rep 12:1497–507.

73. Ko TP, Liao CC, Ku WY, Chak KF, Yuan HS. 1999. The crystal structure of the DNase domain of colicin E7 in complex with its inhibitor Im7 protein. Structure 7:91–102.

74. Kühlmann UC, Pommer AJ, Moore GR, James R, Kleanthous C. 2000. Specificity in protein-protein interactions: the structural basis for dual recognition in endonuclease colicin-immunity protein complexes. J Mol Biol 301:1163–78.

75. Kleanthous C, Walker D. 2001. Immunity proteins: enzyme inhibitors that avoid the active site. Trends Biochem Sci 26:624–31.

76. Morse RP, Willett JL, Johnson PM, Zheng J, Credali A, Iniguez A, Nowick JS, Hayes CS, Goulding CW. 2015. Diversification of beta-Augmentation Interactions between CDI Toxin/Immunity Proteins. J Mol Biol 427:3766–84.

77. Baba T, Ara T, Hasegawa M, Takai Y, Okumura Y, Baba M, Datsenko KA, Tomita M, Wanner BL, Mori H. 2006. Construction of *Escherichia coli* K-12 in-frame, single-gene knockout mutants: the Keio collection. Mol Syst Biol 2:2006 0008.

78. Jones AM, Garza-Sanchez F, So J, Hayes CS, Low DA. 2017. Activation of contact-dependent antibacterial tRNase toxins by translation elongation factors. Proc Natl Acad Sci U S A 114:E1951–E1957.

79. Ferrieres L, Hemery G, Nham T, Guerout AM, Mazel D, Beloin C, Ghigo JM. 2010. Silent mischief: bacteriophage Mu insertions contaminate products of *Escherichia coli* random mutagenesis performed using suicidal transposon delivery plasmids mobilized by broad-host-range RP4 conjugative machinery. J Bacteriol 192:6418–27.

80. Cherepanov PP, Wackernagel W. 1995. Gene disruption in *Escherichia coli*: TcR and KmR cassettes with the option of Flp-catalyzed excision of the antibiotic-resistance determinant. Gene 158:9–14.

81. Jensen SJ, Ruhe ZC, Williams AF, Nhan DQ, Garza-Sánchez F, Low DA, Hayes CS. 2023. Paradoxical Activation of a Type VI Secretion System Phospholipase Effector by Its Cognate Immunity Protein. J Bacteriol 205:e0011323.

82. Hayes CS, Bose B, Sauer RT. 2002. Proline residues at the C terminus of nascent chains induce SsrA tagging during translation termination. J Biol Chem 277:33825–32.

83. Edwards RA, Keller LH, Schifferli DM. 1998. Improved allelic exchange vectors and their use to analyze 987P fimbria gene expression. Gene 207:149–57.

84. Cardona ST, Valvano MA. 2005. An expression vector containing a rhamnose-inducible promoter provides tightly regulated gene expression in *Burkholderia cenocepacia*. Plasmid 54:219–28.

85. Holberger LE, Garza-Sanchez F, Lamoureux J, Low DA, Hayes CS. 2012. A novel family of toxin/antitoxin proteins in *Bacillus* species. FEBS Lett 586:132–6.

86. Whitney JC, Beck CM, Goo YA, Russell AB, Harding BN, De Leon JA, Cunningham DA, Tran BQ, Low DA, Goodlett DR, Hayes CS, Mougous JD. 2014. Genetically distinct pathways guide effector export through the type VI secretion system. Mol Microbiol 92:529–42.

87. Donato SL, Beck CM, Garza-Sánchez F, Jensen SJ, Ruhe ZC, Cunningham DA, Singleton I, Low DA, Hayes CS. 2020. The β-encapsulation cage of rearrangement hotspot (Rhs) effectors is required for type VI secretion. Proc Natl Acad Sci U S A 117:33540–8.

88. Chiang SL, Rubin EJ. 2002. Construction of a mariner-based transposon for epitope-tagging and genomic targeting. Gene 296:179–85.

89. Garza-Sánchez F, Janssen BD, Hayes CS. 2006. Prolyl-tRNA(Pro) in the A-site of SecM-arrested ribosomes inhibits the recruitment of transfer-messenger RNA. J Biol Chem 281:34258–68.

90. Kabsch W. 2010. XDS. Acta Crystallographica Section D: Biological Crystallography D66:125–132.

91. Evans PR, Murshudov GN. 2013. How good are my data and what is the resolution? Acta Crystallographica Section D: Biological Crystallography 69:1204–1214.

92. McCoy AJ, Grosse-Kunstleve RW, Adams PD, Winn MD, Storoni LC, Read RJ. 2007. Phaser crystallographic software. Journal of Applied Crystallography 40:658–674.

93. Terwilliger TC, Adams PD, Read RJ, McCoy AJ, Moriarty NW, Grosse-Kunstleve RW, Afonine PV, Zwart PH, Hung LW. 2009. Decision-making in structure solution using Bayesian estimates of map quality: The PHENIX AutoSol wizard. Acta Crystallographica Section D: Biological Crystallography 65:582–601.

94. Terwilliger TC, Grosse-Kunstleve RW, Afonine PV, Moriarty NW, Zwart PH, Hung LW, Read RJ, Adams PD. 2007. Iterative model building, structure refinement and density modification with the PHENIX AutoBuild wizard. Acta Crystallographica Section D: Biological Crystallography 64:61–69.

95. Afonine PV, Grosse-Kunstleve RW, Echols N, Headd JJ, Moriarty NW, Mustyakimov M, Terwilliger TC, Urzhumtsev A, Zwart PH, Adams PD. 2012. Towards automated crystallographic structure refinement with phenix.refine. Acta Crystallographica Section D: Biological Crystallography 68:352–367.

96. Emsley P, Cowtan K. 2004. Coot: model-building tools for molecular graphics. Acta Crystallogr D Biol Crystallogr 60:2126–2132.

97. Gibson EG, Oviatt AA, Osheroff N. 2020. Two-Dimensional Gel Electrophoresis to Resolve DNA Topoisomers. Methods Mol Biol 2119:15–24.

